# Analysis of tandem mass spectrometry data with CONGA: Combining Open and Narrow searches with Group-wise Analysis

**DOI:** 10.1101/2023.05.02.539167

**Authors:** Jack Freestone, William S. Noble, Uri Keich

## Abstract

Searching tandem mass spectrometry proteomics data against a database is a well-established method for assigning peptide sequences to observed spectra but typically cannot identify peptides harboring unexpected post-translational modifications (PTMs). Open modification searching aims to address this problem by allowing a spectrum to match a peptide even if the spectrum’s precursor mass differs from the peptide mass. However, expanding the search space in this way can lead to a loss in statistical power to detect peptides. We therefore developed a method, called CONGA, that takes into account results from both types of searches—a traditional “narrow window” search and an open modification search—while carrying out rigorous false discovery rate (FDR) control. The result is an algorithm that provides the best of both worlds: the ability to detect unexpected PTMs without a concomitant loss of power to detect unmodified peptides.

## 1. Introductions

Tandem mass spectrometry provides an efficient way to study proteins in a high-throughput fashion but is typically limited in its ability to detect post-translationally modified (PTM) peptides due to the exponentially large search space associated with searching for PTMs. One proposed solution to this problem is to employ “open modification” searching, in which each observed spectrum is compared against peptides whose masses differ—often by hundreds of Daltons—from the observed precursor mass associated with the spectrum ^1^. Open modification searching has become increasingly popular, with one report suggesting that this type of approach can achieve higher statistical power to detect peptides than a traditional “narrow window” database search ^2^.

We recently showed that open searches on their own often produce fewer discoveries than narrow searches applied to the same data ^3^. Only when coupled with a machine learning post-processor such as Percolator ^4^ or PeptideProphet ^5^ do open searches become typically better than narrow window searches. At the same time we provide evidence that both Percolator and PeptideProphet may fail to control the FDR ^3^. Hence, that apparent improvement in power attributed to open modification searching must be taken with a grain of salt.

Motivated by these observations, we propose an alternative post-processor that allows us to consistently deliver more discoveries compared to the traditional narrow search. The key idea is to search the observed spectra twice, once using a wide (open) window and once using a narrow window, and then combine the search results in a novel way that allows us to extract information from both searches while rigorously and empirically controlling the FDR. The method, Combining Open and Narrow searches with Group-wise Analysis (CONGA), draws on the same concept of target-decoy competition (TDC) that is commonly used to control the FDR: each real (“target”) peptide is paired with a randomly shuffled or reversed decoy peptide, and each spectrum is searched against the concatenated target-decoy database. Assuming that a false match is equally likely to involve a target or a decoy peptide, the optimal matches to the decoy peptides allow us to estimate the number of false discoveries and hence control the FDR. In CONGA’s case this step essentially involves a competition scheme that produces a filtered list of peptides, which are then grouped according to the difference between the observed precursor mass and the peptide mass. This grouping is then followed by our FDR analysis, which reports the target peptides exceeding each group threshold while controlling the overall FDR.

We provide empirical evidence for CONGA’s strong performance. First, we verify CONGA’s FDR control using an entrapment experiment. This experiment uses matches to an irrelevant set of peptide sequences, called the entrapment sequences, to essentially calculate the number of incorrect peptide matches in the reported list of peptide discoveries. We then show that CONGA achieves better power to detect peptides than either a narrow or open modification search, and that its power is comparable with that of Percolator. We also demonstrate that CONGA’s approach allows us to detect and better utilize chimeric spectra, because CONGA’s group-wise analysis considers multiple matching peptides for each spectrum while still controlling the overall FDR.

An open source Python implementation of CONGA can be found at https://github.com/freejstone/CONGA.

## 2 Approach

Motivated in part by our doubts about FDR control in existing post-processors ^3^, we developed CONGA. The algorithm takes as input two sets of search results—the top PSM for each spectrum from a narrow search against a concatenated target-decoy database, and the top *n*_*t*_ PSMs for each spectrum from an open search against the same concatenated database. CONGA also requires the target-decoy peptide pairs that make up the concatenated database. The algorithm then proceeds through four main steps, which are summarized in Figure 1 and below, with further details provided in Section 3 (The CONGA algorithm):

**Figure 1:**
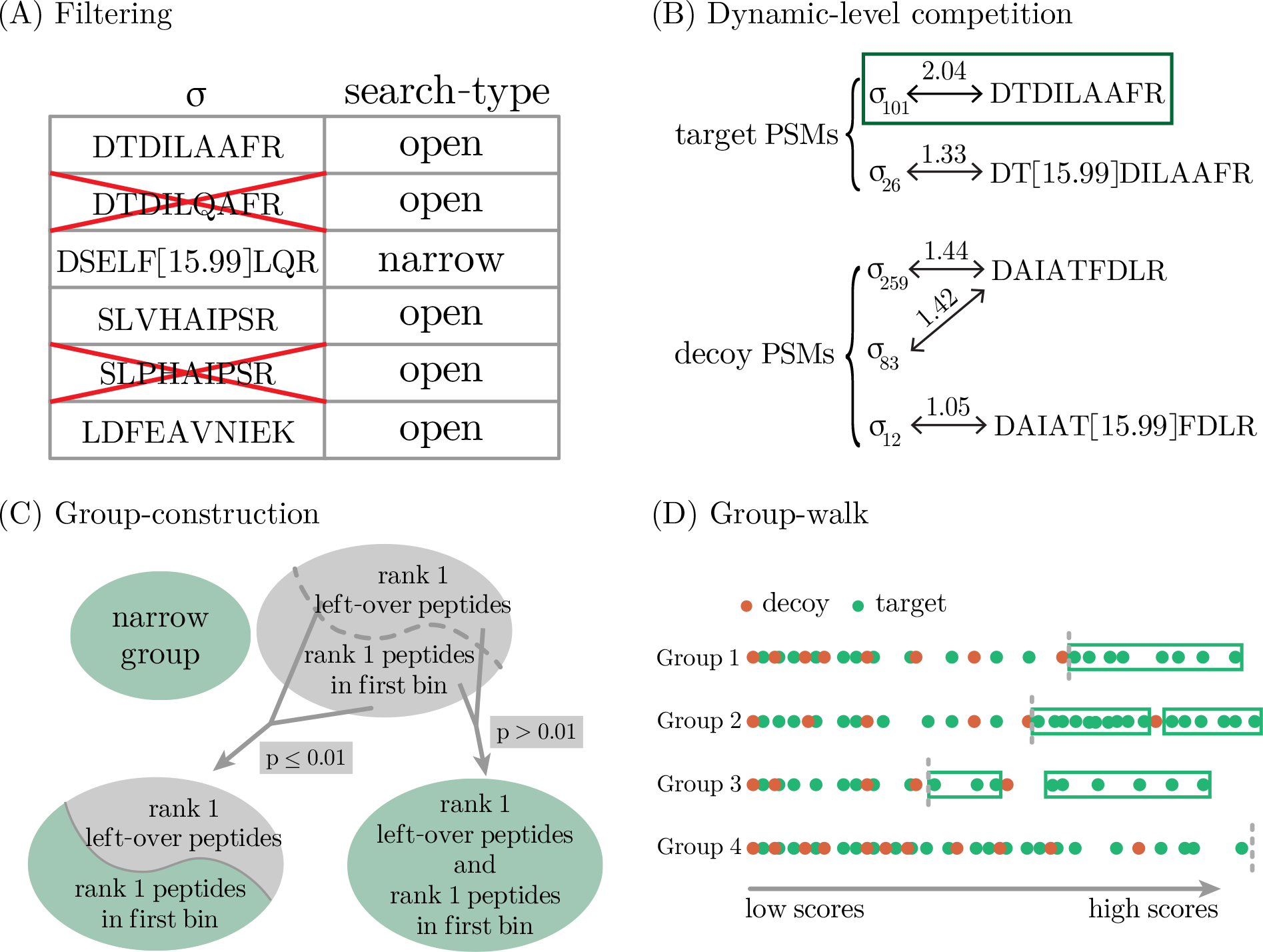
Schematics of CONGA. (A) For each experimental spectrum *σ*, some sub-optimal PSMs need to be weeded out because they involve peptides whose theoretical spectra share a non trivial number of peaks with higher scoring matched peptides. (B) The best scoring peptide among all PSMs that have the same unmodified stem form, along with their decoys, is then selected. (C) Partitioning into groups: the (representative) peptides are first divided into a static “narrow group”, and a second “open group” that initially contains all other peptides. We next adaptively split and merge sub-groups of the open group starting with comparing the winning scores of a “candidate set”, consisting of all rank 1 peptides with the most commonly occurring mass-difference (labeled as the *first bin*), with the winning scores of the remaining rank 1 peptides of the open group. If the p-value of the Kolmogorov-Smirnov test is smaller than 0.01, a new group is created from the candidate set, else a new group is created from all of the rank 1 peptides of the open group (see Group construction section for further details). (D) Once the groups are defined, we employ the Group-walk algorithm which iteratively updates a vector of group-wise thresholds until the estimated FDR from the combined peptides to the right of each threshold is less than *α*. The target peptides above each group-threshold (indicated by the rectangles) are then reported.

1. CONGA’s first step weeds out potentially problematic PSMs. These PSMs arise because CONGA considers more than one PSM per spectrum, allowing the algorithm to properly handle chimeric spectra generated by more than one peptide. The challenge is that some PSMs might “hitchhike” and score highly if the theoretical spectrum of an incorrectly matched peptide closely resembles that of the true, generating peptide. To filter out these PSMs, CONGA first combines the list of narrow- and open-search PSMs. Then for each spectrum, it removes sub-optimal PSMs with peptides that are similar to higher scoring peptides (Figure 1A).
2. CONGA’s second step introduces “dynamic-level competition,” which is designed to help TDC overcome some inherent shortcomings of decoy construction. Specifically, TDC relies on the assumption that a false discovery—in our case a peptide that won its competition but is not in the sample—is equally likely to be a target or a decoy win. However, when variable modifications are specified the database will often contain several clusters of highly similar peptides corresponding to variable modifications of the same unmodified (“stem”) form of a single peptide. For example, consider a spectrum that was generated by P[10]EPTIDE but because of random noise is optimally matched to PEP[10]TIDE. In this case, the canonically-created decoys cannot offer real competition to such a mismatch. Indeed, typically, the theoretical spectra of P[10]EPTIDE and any of its cluster-neighbors, including PEP[10]TIDE, would share significantly more peaks than P[10]EPTIDE and its “closest” decoy would. To address this problem, CONGA dynamically adjusts the level at which TDC takes place. When no variable modifications are specified, the competition is done at the peptide level by taking the best scoring peptide between each pair of target and decoy peptides as described in ^6^. When variable modifications are specified, the competition is done at the stem level. Specifically, CONGA first defines the representative peptides for each stem form as the highest scoring peptide with that stem form. It then competes each pair of target and decoy representative peptides by taking the best scoring one of each pair.
3. CONGA’s third step is the essential one of dividing the list of stem peptides into groups. This fairly involved process, detailed in Section 3 (Group construction), defines the groups based on characteristics of the maximally-matched PSMs associated with the stem peptides including: (a) whether the PSM appeared in the narrow search file (“narrow group”), (b) the frequency of the difference between the precursor mass and the peptide mass (the delta mass), and (c) for PSMs that originate from the open-search file, their rank among all the PSMs of the corresponding spectrum. While the narrow group remains static, the remaining peptides are adaptively split and merged into groups based on applications of the Kolmogorov-Smirnov test to gauge the difference in the winning peptides score distributions between the two currently considered groups.
4. Finally, this grouping is then followed by Group-walk, which works by constructing separate thresholds for each group and subsequently reporting the target peptides above each group threshold. This procedure guarantees theoretical control of the overall FDR while taking advantage of the different characteristics of the groups ^7^. In doing so, CONGA gets around the problem of how to otherwise reward the more popular/reasonable mass modifications. For example, intuitively we should prefer a narrow search PSM over an open search PSM that has a mass modification that appears only once. Rather than trying to adjust the score function, CONGA simply places each PSM in a separate group so their raw scores are never competing against one another.

After reporting its FDR-controlled list of discoveries, CONGA employs two optional steps to further assist the user. The first is complementary to CONGA’s dynamic level competition which is designed with rigorous FDR control in mind. However, regardless of whether searching with variable or static modifications, CONGA’s unique type of competition necessarily leads to some loss of information. For example, particularly when searching without any specified variable modifications, the user might still be interested to learn which unexpected mass modifications of a particular peptide have support in the data. This information is not directly available from CONGA’s FDR-controlled list of discovered peptides. Therefore, CONGA optionally augments this FDR-controlled list by reporting sufficiently high scoring PSMs that are related to the peptides in this list.

In a second optional step, CONGA uses the pyAscore module ^8^ to assist with the localization of modifications in CONGA’s augmented list of PSMs.

## 3 Methods

### The CONGA algorithm

#### Filtering neighbors

CONGA’s first step focuses on removing neighbors: a peptide is considered a *neighbor* of another peptide if the two corresponding theoretical spectra share a non-trivial proportion of peaks. Neighbors pose a problem, particularly if, as in CONGA’s case, we consider more than the top match to a spectrum *σ*: if a peptide offers a good match to *σ* then it is likely that so do all its neighbors.

Supplementary Algorithm S2 describes CONGA’s filtering process. Its input is the list of peptides corresponding to the top peptide-spectrum matches (PSMs) associated with each spectrum. Specifically, associated with each spectrum *σ*are *k*^*σ*^≤ *n*_*t*_ + 1 unique PSMs, of which ≤ *n*_*t*_ are the top PSMs from the open search, and ≤ 1 from the narrow search (we remove an open PSM if it happens to coincide with the narrow PSM). Sorting those *k*^*σ*^peptides by decreasing PSM scores, CONGA sequentially filters out peptides that share more than a fraction of *τ* peaks with a higher ranking peptide (we used *τ* = 0.05, a choice we explain in the Discussion section). Following Lin et al. ^9^, we define *t*_12_, the similarity between two peptides *π*_1_ and *π*_2_, as the proportion of shared *b*- and *y*-ions peaks:

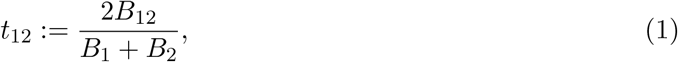

where *B*_*i*_ is the number of charged *b*- and *y*-ions associated with *π*_*i*_ between 200 *m/z* and 3000 *m/z*, and *B*_12_ is the number of such shared ions. If the presumed charge of *π*_*i*_ is +1 or +2, then *B*_*i*_ considers only singly charged *b*- and *y*-ions, and if the charge is +3 or more, then *B*_*i*_ considers both singly and doubly charged ions.

Comet’s E-value ^10^ and Tide’s Tailor score ^11^ normalization of the XCorr score will generally differ between the open and narrow searches. Therefore, to ensure a fair comparison when sorting the combined list of *k*^*σ*^ peptides we use the PSMs’ XCorr score. The PSMs that are not filtered out are then reverted back to their original E-value or Tailor score.

#### Dynamic Competition

CONGA’s second step performs the competition that defines the target/decoy labels on which TDC relies on. Note that this is the second competition, with the first one taking place at the PSM level even before CONGA starts: each spectrum is searched for its best matches in the concatenated target-decoy database. CONGA then filters the PSMs as described above, and the remaining PSMs are used to assign a (peptide-level) score to each target and decoy peptide: the score of the maximal PSM associated with that peptide (or the smallest possible score if no such PSM exists).

The second level of competition differs depending on whether the user specified variable modifications during the search. Importantly, each decoy peptide is constructed by reversing or randomly shuffling the target peptide, and can thus be paired with the target peptide that generated it. When no variable modifications are specified, the second target-decoy competition takes place at the peptide level: each target peptide is compared with its corresponding decoy, and only the higher scoring of the two is kept. On the other hand, when variable modifications are specified, CONGA clusters together all peptides that share a common stem form (the unmodified peptide sequence). Each cluster is represented by its maximally scoring peptide. In this case, the second target-decoy competition takes place at the representative, rather than the peptide level: each target representative is compared with its corresponding decoy representative. Note that, as alluded to above, regardless of whether variable modifications are considered, CONGA optionally produces an auxiliary list of discoveries that includes all sufficiently high scoring PSMs with distinct unaccounted-for mass modifications (details below).

After the second-level competition takes place, each winning peptide is assigned five values: its winning score (*W*_*i*_), the target/decoy label (*L*_*i*_ = 1 if the winner is a target object), a narrow-search indicator (*N*_*i*_ = 1 if the maximal PSM of the winner came from a narrow search), the mass difference between the precursor mass and the peptide (*δ*_*i*_), and finally the spectrum-specific rank *R*_*i*_ of the winning peptide among the top PSMs reported for that spectrum in the original search file. Because we only keep the top PSM per spectrum in the narrow search, *R*_*i*_ = 1 if *N*_*i*_ = 1, but at any rate CONGA only further considers peptides/representatives for which *R*_*i*_ ≤ 2.

#### Group construction

CONGA’s third step involves aggregating peptides into groups. The motivation for this grouping step is that such groups, when they reflect inherent properties of the data, can lead to a boost in statistical power. For example, we expect the narrow peptides (*N*_*i*_ = 1) to be distinct from the open ones; hence, those peptides form their own “narrow group”. The other groups, made up of the open peptides (*N*_*i*_ = 0), are essentially defined based on the mass differences *δ*_*i*_. As outlined next, and detailed in Supplementary Algorithm S3, CONGA accomplishes this grouping in two phases, in which it dynamically splits and merges the open peptides according to *δ*_*i*_, in an attempt to define distinct groups in terms of their winning scores distribution. Note that throughout this section we refer only to “peptides,” with the understanding that we mean representatives when using variable modifications.

With the narrow group defined, CONGA first partitions the open peptides with rank *R*_*i*_ = 1 into a set of disjoint bins according to their associated mass differences, *δ*_*i*_. Specifically, each bin is centered at *kλ*, an integer multiple of *λ* := 1.0005079*/*4, and it contains all peptides with *δ*_*i*_ ∈ (*kλ* − *λ/*2, *kλ* + *λ/*2]. We then rank the bins in terms of their occupancy rate from largest to smallest, and we denote by *b*_*i*_ the rank of the bin containing the *i*th considered peptide.

In practice, CONGA cannot make efficient use of a group which is too small; hence, it is better to merge such a group with another one. CONGA’s minimal group size is set to 2*K* (80 by default), where *K* is the window size used by Group-walk, as explained in the next section. By merging any bin that is smaller than this threshold with the subsequently ranked bin, we can assume without loss of generality that every bin, except possibly the lowest ranked one, clears the minimal group size threshold.

Starting with the top ranked bin (i.e., the most common mass difference), CONGA sequentially defines a candidate set *C* as the set of peptides populating the highest ranked bin that is yet to be processed. Defining the current left-over set, *L*, as the set of peptides in all lower ranked bins, CONGA applies the Kolmogorov-Smirnov (KS) test to gauge the similarity between the winning scores of (i) *C* and *L*, and (ii) *C* and each group of open peptides previously defined in this sequential process. For example, when considering the 3rd ranked bin this would typically generate 1 + 2 = 3 p-values, one for each application of the test.

If all the p-values are less than or equal to a pre-specified threshold *β* := 0.01, then the winning scores in *C* are deemed distinct enough to merit defining *C* as a new group, and the process moves on to the next candidate set. Otherwise, if the p-value of the test comparing *C* and *L* is ≤ *β* but the p-values of the similarity tests between *C* and some of the previously defined groups are *> β*, then *C* is deemed to be insufficiently distinct from any one of those groups. In that case, one of those groups for which the KS p-value is *> β* is selected at random, *C* is merged with it, and the process moves on to the next candidate set. Otherwise, the p-value of the similarity test between *C* and *L* is *> β* so *C* and *L* are joined into a new group, and this iterative phase of the group construction ends.

If the process did not terminate before *C* coincides with the lowest ranked bin, then at that point *C* is added to the most recently defined group. Note that in practice the winning PSM scores often have ties, which implies that the Kolmogorov Smirnov p-values are only approximate. However, this behavior is not overly concerning because we are only using the tests here as a heuristic group-merging criterion rather than for hypothesis testing per se.

Finally, CONGA groups together all open peptides with rank *R*_*i*_ = 2 and a mass difference *δ*_*i*_ that would place it in one of the top ranking bins that were processed in the previous phase (excluding the terminating step). If this group is larger than the 2*K* threshold then it is defined as the last group; otherwise, it is merged with the most recently constructed group (which is usually the final candidate set *C* and the left-over set *L*).

#### Group-walk

After the groups are constructed, CONGA applies the Group-walk procedure ^7^, which is added here for convenient reference as Supplementary Algorithm S4. Briefly, Group-walk first orders the peptides (representatives) within each group *g* ∈ {1, …, *n*_*g*_} in increasing winning scores order, so that with *n*_*g*_ being the number of groups, 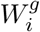 the score of the *i*th peptide in group *g*, and *m*_*g*_ the number of peptides in group *g*, 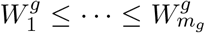 (note that this ordering is reversed to the usual way TDC is formulated).

CONGA then initializes a *frontier* vector 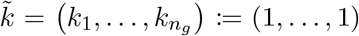, which delineates the “active set” of peptides that the front has not yet passed over: all peptides for which the (group *g*) index *i* satisfies *i* ≥ *k*_*g*_ (so 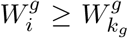). The peptides that are not in the active set cannot be discovered (the front passed over them), but as explained below, they are still useful in determining how to advance the front.

Group-walk advances the front one coordinate at a time, and following each such advance it estimates the FDR among the target peptides in the active set as the number of decoys in this set (plus 1) divided by the number of target peptides in this set. If this estimated FDR is ≤ *α* then Group-walk stops and reports all target peptides in the active set. Otherwise, Group-walk selects a group *g* for which it will advance the front by one peptide: *k*_*g*_ := *k*_*g*_ + 1.

Initially, the group that is advanced is selected sequentially: Group-walk rotates through the groups. However, once the frontier reaches 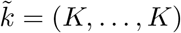, where *K* is the “peek-back” window size (*K* = 40 by default), the decision of which group to advance is made based on the last *K* observed peptides in each group. Specifically, Group-walk advances the group for which the number of decoy wins among the last *K* peptides is maximized.

#### Auxiliary list of discoveries

CONGA optionally complements it list of FDR-controlled discoveries by producing an auxiliary list of discoveries that includes all sufficiently high scoring PSMs with distinct unaccounted-for mass modifications explained next (Supplementary Algorithm S6).

If no variable modifications are specified, then for each peptide in its FDR-controlled list, CONGA reports any PSM involving the same peptide that (a) is sufficiently high scoring, i.e., it is a top-ranked narrow, or a top-2 open PSM (*R*_*i*_ ≤ 2), and (b) is the highest scoring of all the PSMs with the same peptide that fall into the same mass modification bin. The mass bins are defined by a greedy clustering algorithm (Supplementary Algorithm S5) that takes into account the specified isolation window. This clustering is necessary to avoid overwhelming the user with PSMs that share the same peptide and have very similar precursor masses. The augmented list is further supported with a label for each PSM which indicates whether the PSM scored above its CONGA-assigned group-specific threshold, giving the user additional confidence in those PSMs.

When variable modifications are specified, the process is slightly different. Specifically, for each representative peptide in the FDR-controlled list, CONGA reports any PSM that (a) is sufficiently high scoring as defined above, (b) its peptide shares the same stem as the representative’s, and (c) is the highest scoring among all PSMs with the same variant of the stem form that fall into the same mass modification bin. In this case the greedily-clustered bins are defined relative to the unmodified (stem) form.

#### Localization with pyAscore

CONGA optionally uses the pyAscore module ^8^ to assist in the localization of modifications in CONGA’s augmented list of PSMs. Specifically, for each PSM with a delta mass *δ*_*i*_ outside the specified “narrow search” window, pyAscore enumerates all possible localizations of *δ*_*i*_ and chooses the highest scoring one. Furthermore, if the user provides a list of anticipated modifications, then pyAscore will also attempt to localize any such modification that is sufficiently close to the observed *δ*_*i*_. Finally, CONGA will report for any such PSM the single highest scoring of all those localizations, along with a label indicating whether or not that optimal localization scored higher than the null modification (that is, when no localization is considered).

### Searches

#### Entrapment run searches

Entrapment runs for CONGA were performed using data taken from the standard protein mix database, ISB18 ^12^. We used the nine .ms2 spectrum files taken from Lin et al. ^6^ which were originally sourced from the Mix 7 spectrum files downloaded from https://regis-web.systemsbiology.net/PublicDatasets/18_Mix/Mix_7/ORBITRAP/ (this excluded the tenth, smallest spectrum file). These .ms2 spectrum files were subsequently used for Tide. For compatibility with MSFragger, we directly downloaded the same nine spectrum files in ThermoRaw format and converted them to a single combined, as well as separate, .mzML files using MSConvert 3.0.22314 with the vendor peak-picking filter using the default settings.

The in-sample database contained peptides from the original 18 proteins plus 30 additional hitchhiker proteins believed to be present at lower levels, which we downloaded from https://regis-web.systemsbiology.net/PublicDatasets/database. We used the castor plant proteome as the entrapment database as in Lin et al. ^6^. Tide-index was used to digest the in-sample and entrapment database using trypsin, for Tide, and strict-tryspin (specified with `trypsin/p`), for MSFragger. For each digest rule, four random subsets of the in-sample peptides of varying size, plus the entrapment peptides, were used to create a combined target peptide database for which 20 randomly shuffled decoy indices were created, again using Tide-index. Param-medic ^13^ was used to determine any variable modifications, though none were detected. Any entrapment peptide that was identical (up to any leucine/isoleucine substitution) to an ISB18 peptide was removed from the entrapment database. Table 1 lists the resulting number of in-sample and entrapment peptides in each combined target peptide database, along with their in-sample-to-entrapment ratio.

**Table 1:**
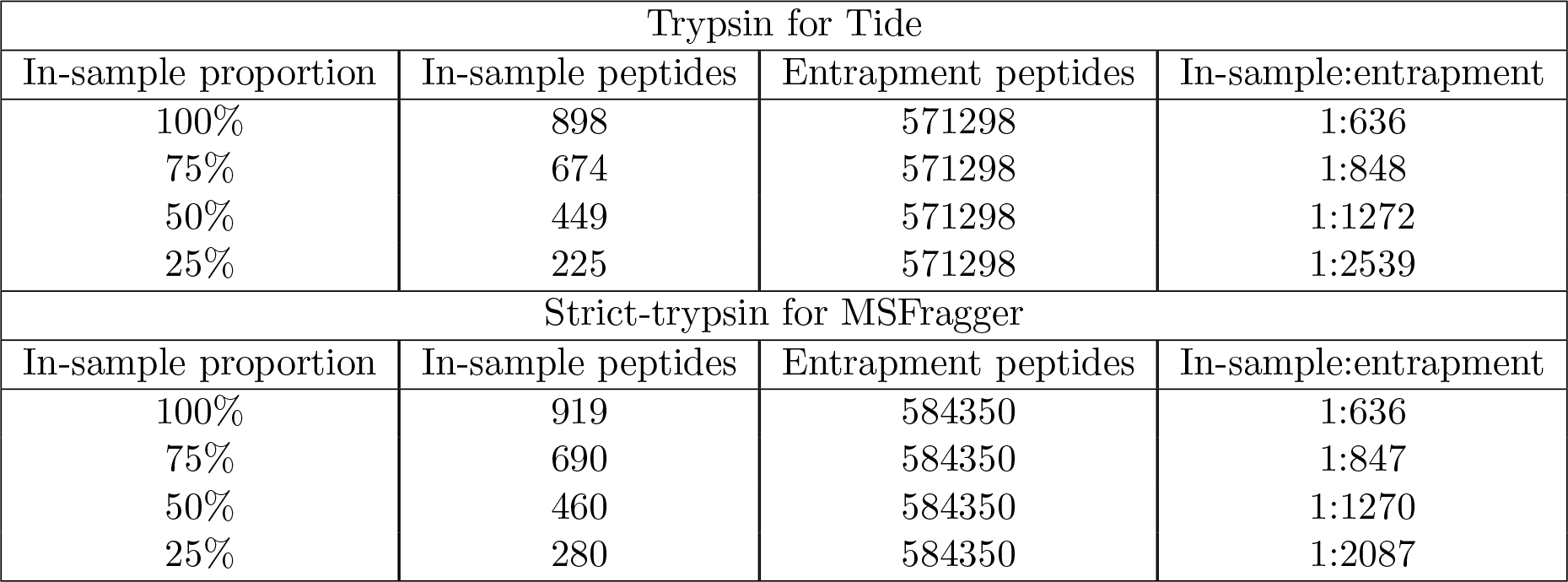
The number of in-sample target peptides and entrapment-peptides for each combined target peptide database used in the entrapment runs. The table reports the number of in-sample target peptides and entrapment target peptides for each of the four combined target databases, for each digest rule. The first column reports the proportion of the total in-sample target peptides used in the combined target database, the second is the number of these in-sample peptides used, the third column is the number of entrapment peptides, and the last column is the ratio of in-sample peptides to entrapment peptides.

Tide-search was used to search the nine .`ms2` spectrum files against each of the combined target and decoy databases prepared using trypsin with the following settings: for narrow searches we used `--auto-precursor-window warn --auto-mz-bin-width warn --use-tailor-calibration T --concat T`, and for open searches we used `--auto-mz-bin-width warn --precursor-window-type mass --precursor-window 100 --top-match 5 --concat T --use -tailor-calibration` T. MSFragger v3.3 was used to search the combined .mzML and the separate .mzML spectrum files against each of the combined target and decoy databases prepared using strict-trypsin. Because we used tide-index to prepare these databases, we need to use the following settings to ensure MSFragger has the same digestion parameters. For narrow searches we used `allowed_missed_cleavage = 0, digest_min_length = 6, digest_max_length = 50, digest_mass_range = 200.0 7200.0, allowed_missed_cleavage = 0` with all other options set to default, and for open searches we used `localize_delta_mass = 0, allowed_missed_cleavage = 0, digest_min_length = 6, digest_- max_length = 50, digest_mass_range = 200.0 7200.0, allowed_missed_cleavage = 0, output_report_topN = 5`, with all other option set to default. No variable modifications were set, so CONGA’s analysis was done at the peptide level. Note we used MSFragger v3.3 at the time, as the subsequent version at the time (v3.4) did not allow for `output_report_topN > 1` for DDA data. We used Tide in Crux v4.1.decd99ff.

#### PRIDE-20 searches

We downloaded 20 high-resolution spectrum files from the Proteomics Identifications Database, PRIDE ^14^. Seven of these spectrum files were taken from Freestone et al. ^7^ and were originally obtained by randomly selecting a spectrum file from randomly selected PRIDE projects (submitted no earlier than 2018). The remaining 13 spectrum files were similarly obtained by randomly selecting a single spectrum file from a randomly selected PRIDE project (submitted no earlier than 2019). The sampling was constrained to generate a collection of 10 spectrum files that had modifications and 10 spectrum files for which no modifications were detected, as determined in both cases by Param-medic ^13^. The protein FASTA database files were also downloaded from the associated PRIDE projects or, in the case of human data, the UniProt database UP000005640 was used (downloaded 9/11/2021). Table 2 reports the list of the 20 spectrum files and PRIDE projects used.

**Table 2:**
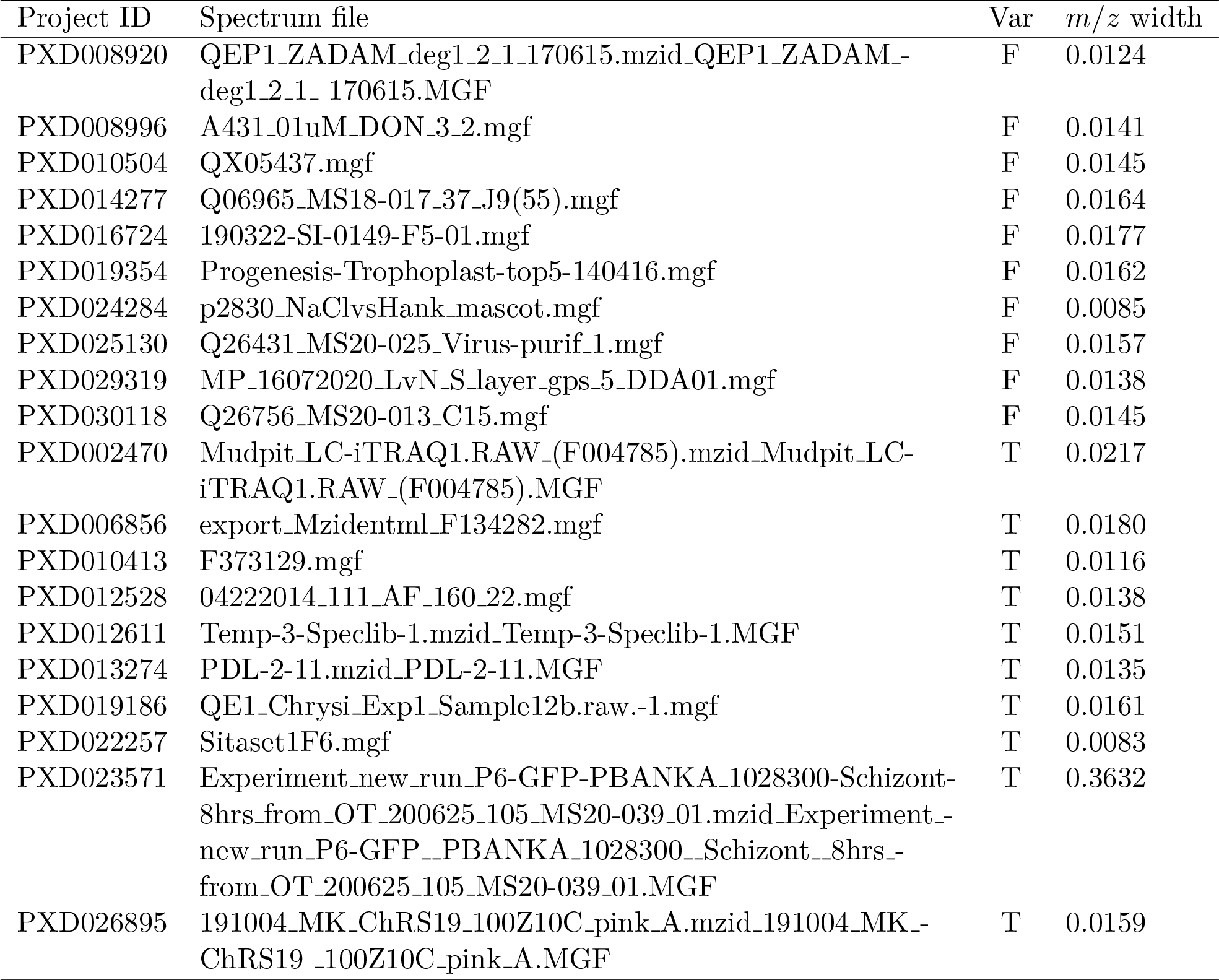
The PRIDE-20 data set. The list of 20 spectrum files used in the PRIDE-20 data set and their associated project IDs. The “Var” column indicates whether the data includes variable modifications. The “*m/z* width” column reports the fragment *m/z tolerances determined by Param-medic* ^*15*^.

| Project ID | Spectrum file | Var | $m/z$ width |
| --- | --- | --- | --- |
| PXD008920 | QEP1_ZADAM_deg1_2_1_170615.mzid_QEP1_ZADAM_-deg1_2_1_170615.MGF | F | 0.0124 |
| PXD008996 | A431_01uM_DON_3_2.mgf | F | 0.0141 |
| PXD010504 | QX05437.mgf | F | 0.0145 |
| PXD014277 | Q06965_MS18-017_37_J9(55).mgf | F | 0.0164 |
| PXD016724 | 190322-SI-0149-F5-01.mgf | F | 0.0177 |
| PXD019354 | Progenesis-Trophoplast-top5-140416.mgf | F | 0.0162 |
| PXD024284 | p2830_NaClvsHank_mascot.mgf | F | 0.0085 |
| PXD025130 | Q26431_MS20-025_Virus-purif_1.mgf | F | 0.0157 |
| PXD029319 | MP_16072020_LvN_S_layer_gps_5_DDA01.mgf | F | 0.0138 |
| PXD030118 | Q26756_MS20-013_C15.mgf | F | 0.0145 |
| PXD002470 | Mudpit_LC-iTRAQ1.RAW_(F004785).mzid_Mudpit_LC-iTRAQ1.RAW_(F004785).MGF | T | 0.0217 |
| PXD006856 | export_Mzidentml_F134282.mgf | T | 0.0180 |
| PXD010413 | F373129.mgf | T | 0.0116 |
| PXD012528 | 04222014_111_AF_160_22.mgf | T | 0.0138 |
| PXD012611 | Temp-3-Speclib-1.mzid_Temp-3-Speclib-1.MGF | T | 0.0151 |
| PXD013274 | PDL-2-11.mzid_PDL-2-11.MGF | T | 0.0135 |
| PXD019186 | QE1_Chrysi_Exp1_Sample12b.raw.-1.mgf | T | 0.0161 |
| PXD022257 | Sitaset1F6.mgf | T | 0.0083 |
| PXD023571 | Experiment_new_run_P6-GFP-PBANKA_1028300-Schizont-8hrs_from_OT_200625_105_MS20-039_01.mzid_Experiment_new_run_P6-GFP_PBANKA_1028300_Schizont_8hrs_from_OT_200625_105_MS20-039_01.MGF | T | 0.3632 |
| PXD026895 | 191004_MK_ChRS19_100Z10C_pink_A.mzid_191004_MK_ChRS19_100Z10C_pink_A.MGF | T | 0.0159 |

For each of the PRIDE-20 data sets, we used Tide-index 20 times to create 20 different target-decoy peptide databases using --auto-modifications T to call Param-medic and apply the variable modifications. For narrow searches using Tide, we used the following options --auto-precursor-window warn --auto-mz-bin-width warn --use-tailor-calibration T --concat T and for open searches using Tide we used `--auto-mz-bin-width warn --precursor-window-type mass --precursor-window 100 --use-tailor-calibration T --concat T`.

For searches using Comet, we used the built-in version within Crux. For narrow searches we used the options `--decoy_search 1 --auto_peptide_mass_tolerance warn --allowed_ missed_cleavage 0 --auto_fragment_bin_tol warn --auto_modifications T --peptide _mass units 2` and for open searches we used `--decoy search 1 --peptide_mass_tolerance 100 --allowed_missed - cleavage 0 --auto_fragment_bin_tol warn --auto_modifications T --peptide_mass_units 0`. Comet digests and produces decoy peptides by reversing each target sequence. Hence, only a single target-decoy database was used for each of the PRIDE-20 data sets. PSMs matched to peptides that were identified as both a target and decoy were deleted. In the case of E-values, lower scores indicate a better PSM and so we take the negative of these scores when applying TDC or CONGA.

Lastly, for searches using MSFragger we converted each .`mgf` file into .`mzmL` files using MSCon-vert with default settings. We constructed a decoy protein database by taking each target protein and reversing the position of the amino acids that lie between *K, R* or the *N* - and *C*-terminal, while keeping the position of *K* and *R* fixed. Importantly, the digestion of the target and decoy proteins allow us to pair each resulting target peptide with a corresponding decoy peptide. This is not achievable if we reverse the entire target protein sequence, which is the behavior of many software tools. For narrow searches, we used the following options `allowed_missed_- cleavage = 0, use_topN_peaks = 100` and for open searches we used `output_report_topN = 5, allowed_missed_cleavage = 0, use_topN_peaks = 100`. Variable modifications were set according to those detected by Param-medic using Tide-index. All other options were set to the default settings. Similar to Comet, PSMs matched to peptides that were identified as both a target and decoy were deleted.

#### HEK293 searches

We downloaded the 24 HEK293 spectrum files from https://ftp.pride.ebi.ac.uk/pride/data/archive/2015/06/PXD001468, which were converted to .mzML format using MSConvert with the vendor peak-picking filter using the default settings. We used the UniProt database UP000005640 (downloaded 18/05/2022). For the results that compare CONGA to other methods, we followed the exact same method for searching with Tide and Comet with the 20 PRIDE-20 spectrum files as outlined in the previous section, PRIDE-20 searches. For MSFragger searches we prepared a combined target-decoy protein database similarly described in section, PRIDE-20 searches, by taking each target protein and reversing the position of the amino acids that lie between *K, R* or the *N* - and *C*-terminal, while keeping the position of *K* and *R* fixed, again deleting peptides that were identified as both a target and decoy. We ran narrow MSFragger searches for each of the 24 spectrum files using the settings `precursor_mass_lower = -100, precursor_- mass upper = 100, allowed_missed_cleavage = 1, use_topN_peaks = 100` and open searches using the settings `precursor_mass_lower = -500, precursor_mass_upper = 500, precursor_- mass_units = 0, localize_delta_mass = 0, allowed_missed_cleavage = 1, output_report_- topN = 5`. No variable modifications were considered, and all other options were set to default. These settings were used to follow as best we reasonably could the setup used by Kong et al. ^2^.

For some of the experiments in this paper we focused on a single randomly selected HEK293 dataset, `b1924_293T_proteinID_04A_QE3_122212.mzML`, which we refer to as “the representative HEK293 dataset.”

#### Chimera spectra analysis

We used Prosit to validate our co-peptide (i.e., pairs of peptides assigned to a single spectrum) discoveries in the representative HEK293 dataset. For this analysis, we first converted the data to mgf format with MSConvert for easy importing in R. We determined the list of co-peptide discoveries using CONGA at the 1% FDR level, and we filtered this list for co-peptides that were both less than 1 Da away from the precursor mass of the corresponding experimental spectrum. This is because Prosit does not generally support peptides with mass modifications. In addition to these co-peptides, we determined their precursor charge and set the collision energy for each peptide to 35%, as described in Chick et al. ^1^. The peptides, charges, and collision energy were then submitted to Prosit. The resulting output returned the predicted fragment *m/z* locations, the relative intensities and the normalized retention times (iRT).

We then looked to pair each Prosit predicted peak of each discovered co-peptide with a corresponding peak from the chimeric spectrum that was responsible for discovering the co-peptide. Specifically, we searched the spectrum for experimental peaks within a 0.01 *m/z* fragment tolerance of the Prosit peak. If there was zero or one experimental peak within that window, then we paired the Prosit peak with the experimental peak, or with a zero intensity peak if there was no experimental peak within that window. If there was more than one experimental peak within the window, then we removed the Prosit peak from further consideration because we could not uniquely match it to an experimental one. Finally, we transformed the paired experimental intensities into relative intensities, by dividing the experimental intensities by the maximal intensity of all paired experimental peaks defined for the given co-peptide.

To benchmark our co-peptide discoveries, we determined the list of narrow discoveries using TDC at the 1% FDR level, and similarly used Prosit along with the subsequent analysis as described above for the co-peptide discoveries using CONGA.

#### Target-decoy competition

For the TDC procedure, we employ the double competition protocol, “PSM-and-peptide,” as originally described in Lin et al. ^6^ and summarized for convenience below, and in pseudocode in Supplementary Algorithm S1a. For the PSM-level competition, we determine the best peptide-spectrum match to the target and decoy databases separately, and then record the best scoring match out of the two. This is achieved implicitly by taking the top 1 PSM for each spectrum when searching against the concatenated target-decoy database. For the subsequent peptide-level competition, we define a score for each peptide as the maximal PSM score associated with that peptide. Then for each target-decoy pair, we take the peptide with highest peptide score. The resulting peptides are then ordered according to their peptide score from largest to smallest. Denoting the resulting scores as *W*_1_ ≥ · · · ≥ *W*_*n*_, we determine the largest index *k* that the estimated FDR is ≤ the threshold *α*, and report all target peptides up to and including that *k*. More specifically, denoting *L*_*i*_ = 1 as a target peptide and *L*_*i*_ = −1 as a decoy peptide, then all target peptides with rank *i* ≤ *K*_TDC_ are reported, where

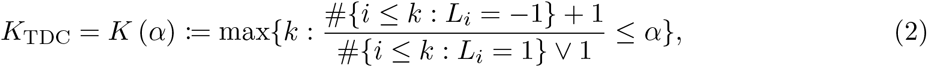

and *x* ∨ *y* := max{*x, y*}.

As described in Section 3 (Dynamic Competition), if variable modifications are specified, then the second, peptide-level competition is replaced with the representative-level one (Supplementary Algorithm S1b).

#### Percolator settings

For each of the Tide searches where Percolator was subsequently used, we constructed .pin files by either specifying `--pin-output T` during Tide-search or by using the `make-pin` utility command in Crux. When creating .pin files from an open search we used the default number of top 5 PSMs which allows for generating the feature `deltLCn` (the difference between the XCorr score of the top 1 PSM and the last ranked PSM). We then kept only the top 1 PSM for each spectrum to overcome the problem of neighbors. For narrow searches, we directly used the top 1 PSM. Lastly, to each PSM in the .`pin` file, we set the `enzInt` feature (the number of internal enzymatic sites) to zero as explained in Danilova et al. ^16^.

For each of the .pin files, we used the following options in the crux percolator command `--only-psms T, --tdc F`. This reports only the PSMs and does not apply the TDC procedure within Percolator since it does not use the double competition protocol, “PSM-and-peptide”. Instead, we apply our TDC procedure as described in the previous section, to the resulting list of PSMs, using the learned percolator score as the PSM score.

## 4 Results

### CONGA successfully controls the FDR using target-decoy competition

TDC established itself as the canonical approach to FDR control in mass spectrometry analysis even before the corresponding mathematical theory was developed independently ^17,18^. That theory is based on the following two assumptions: (a) an incorrect peptide (one that is not present in the sample) that won its target-decoy competition is equally likely to be a target or a decoy peptide, and (b) this happens independently of everything else (the scores of all the peptides, including the considered one, as well as all the target/decoy label of all other peptides).

CONGA relies on an extension of that theory, where the above independence is assumed to hold given the scores as well as the group structure. More precisely, we assume that, conditioned on all the winning (stem) peptide scores (step 2), on CONGA’s group partition (step 3), and on all the target/decoy labels of the peptides that are truly in the sample, the peptides that won their competitions and are not in the sample are independently equally likely to be a target or a decoy win. Under these assumptions Group-walk, and hence CONGA, are guaranteed to control the FDR ^7^.

In practice these theoretical assumptions are naturally met with some approximation; thus, it is important that we test CONGA’s ability to control the FDR using the same entrapment setup in which Percolator and PeptideProphet apparently fail to do so ^3^. Specifically, we applied CONGA to PSMs generated by Tide and MSFragger, searching the same concatenated databases in both narrow and open mode. Reassuringly, the analysis of CONGA’s results in these entrapment experiments (Figure 2) show that the maximal estimated FDR is always below or very close to the corresponding FDR threshold, suggesting that CONGA successfully controls the FDR.

**Figure 2:**
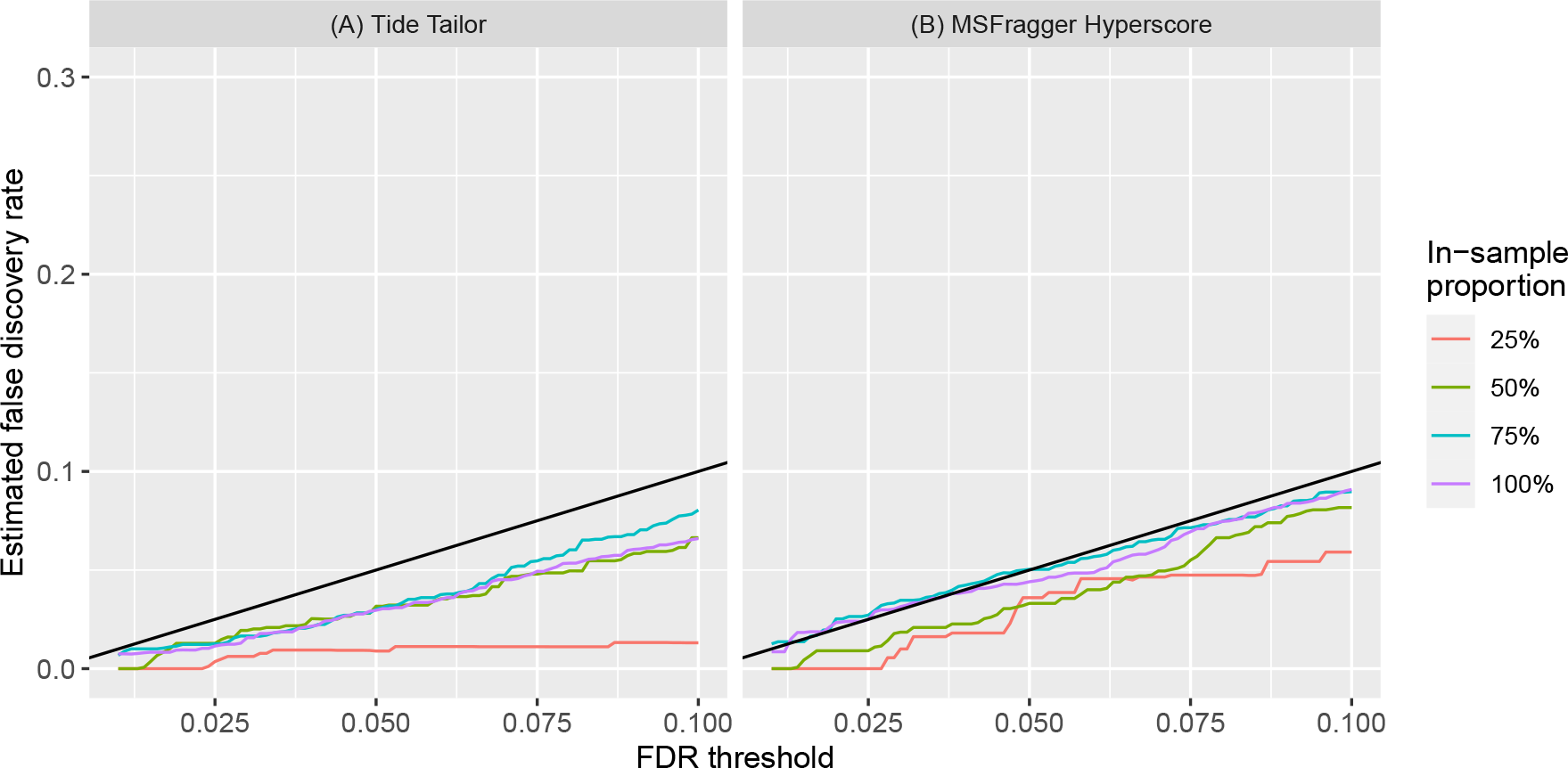
CONGA’s empirical FDR. The estimated FDR when applying CONGA to our entrapment setup using (A) Tide and (B) MSFragger. The FDP is estimated at a range of FDR thresholds ([0.01, 0.1]), and its average over the 20 randomly generated decoys is the empirical FDR. Each solid curve represents the FDR using a different target database constructed with the specified proportion of the in-sample database. The estimated FDR is always below or just above the specified threshold (solid black line), indicating that the FDR is apparently controlled.

### CONGA detects more peptides than narrow or open search

CONGA’s primary goal is to combine the results from narrow and open searches of the same data to yield better statistical power than either of those two searches on its own. Accordingly, we compared the number of peptides detected by CONGA with the number detected by TDC on narrow and open search results. For this analysis, we applied Tide, Comet and MSFragger to our “PRIDE-20” set (Section 3; PRIDE-20 searches): a set of 20 high-resolution spectrum files randomly selected from the Proteomics Identifications Database, PRIDE^14^. TDC was conducted with dynamic-level competition similar to CONGA (Section 3; Target-decoy competition).

The experiments show a consistent improvement in power for CONGA over both narrow and open searching, using all three search engines (Figure 3A–F). Specifically, the median ratio of the number of peptides discovered by CONGA relative to either the narrow or open search is always greater than 1. At 1% FDR, CONGA delivers a median increase over the narrow search of 9.23% for Tide, 21.45% for Comet, 10.95% for MSFragger, and increases over the open search of 18.40%, 17.73%, and 20.37%, respectively. Even at the lower quartiles, CONGA maintains an increase of at least 3% over TDC across all three search engines at both 1% and 5% FDR thresholds. The exact ratios for each of the PRIDE-20 datasets is given in the Supplementary Table S1, and an analogue plot of Figure 3 is given in the Supplement (see Figure S4), which shows the total number of discoveries instead of the ratio of discoveries of CONGA to other methods.

**Figure 3:**
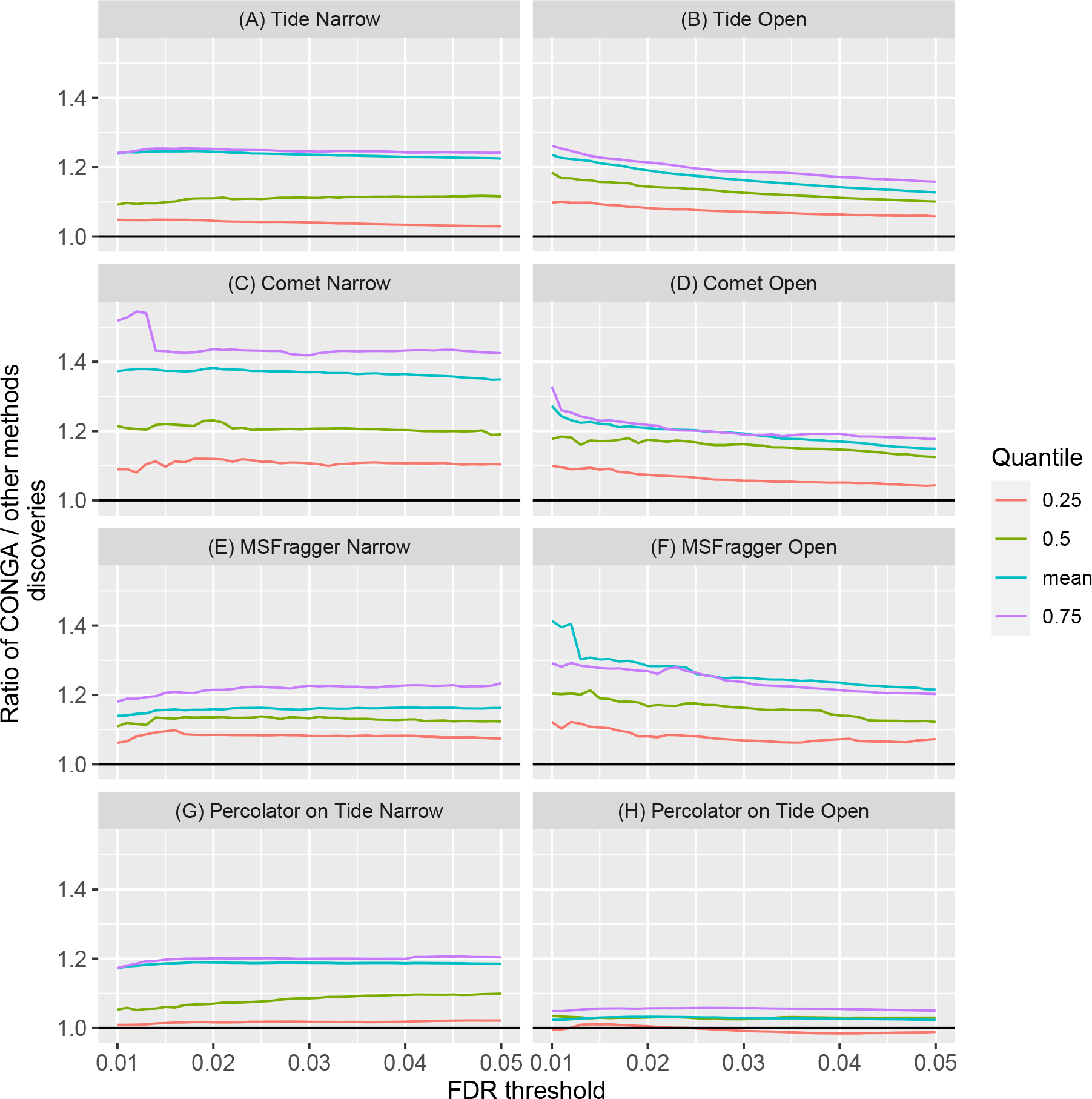
CONGA detects more peptides than narrow or open search. The quartiles and average, taken over the 20 PRIDE-20 datasets, of the mean ratio of CONGA-to-TDC discovered peptides. TDC was applied to (A) Tide search results in narrow mode, (B) Tide search results in open mode, (C) Comet search results in narrow mode, (D) Comet search results in open mode, (E) MSFragger search results in narrow mode and (F) MSFragger search results in open mode. Panels (G) and (H) are similar, but compare CONGA with Percolator applied to the Tide narrow and open search results, respectively. For results that used Tide the mean ratio is taken with respect to each dataset’s 20 decoys, while Comet’s and MSFragger’s searches used a single reversed set of decoys.

Perhaps the key challenge associated with open searching is that the large number of candidate peptides per spectrum leads to a loss of power relative to narrow searching. This occurs when a correctly matched peptide in the narrow search gets out-competed by a candidate from the wide window with a randomly high score. We therefore examined how many peptides discovered in the narrow search are lost in the corresponding list of CONGA discoveries. Specifically, we calculated the number of peptides discovered by Tide with a narrow search that were missing from those reported by CONGA (Supplementary Figure S2(A)). At 1% FDR, the median and mean number of discoveries lost were 85 and 112 peptides respectively. While not completely negligible, this loss is clearly more than offset by CONGA’s gains from the open search. Indeed, the median and mean number of discoveries at the 1% FDR made outside the narrow-window precursor tolerance was 519 and 772 peptides, respectively (Supplementary Figure S2(B)). Moreover, this loss is comparatively far better than the loss when using open-TDC. We calculated the number of narrow-TDC discoveries that are also discovered by CONGA minus the number of narrow-TDC discoveries that are also discovered by open-TDC (Supplementary Figure S3(A)). The median and mean difference at the 1% FDR were 331 and 556 peptides, respectively. CONGA is thus able to retain a large number of peptide discoveries found using a narrow-search.

We then compared CONGA’s ability to detect modified peptides with open-TDC. In particular, we considered the difference in the number of discoveries outside the narrow window precursor tolerance made by CONGA and open-TDC (Supplementary Figure S3(B)). CONGA retains a majority of these discoveries at the 1% FDR level, with a median loss of 12 peptides overall and a mean gain of 69 peptides. Similarly, we looked at the difference in the number of discoveries with delta-masses that coincide with oxidation (Supplementary Figure S3(C)). At 1% FDR, CONGA only has a median loss of 1 peptide discovery and a mean gain of 18 peptide discoveries. The large discrepancies between the median and mean is driven by a handful of datasets (PXD019186, PXD026895, PXD016724, PXD030118) that appear to have an enriched number of modified peptides.

A key remaining question is how CONGA compares to a post-processor such as Percolator. Carrying out this comparison is challenging because, as pointed out in Freestone et al. ^3^, Percolator’s FDR control seems to be liberally biased. Thus, it is particularly interesting that, overall, our experiments suggest that CONGA marginally edges out Percolator when both are applied to the PRIDE-20 datasets (Figure 3G–H). Specifically, both post-processors use Tide open and narrow searches, where Percolator can only use one type of search at a time but is able to use significantly more features than CONGA can. For both open and narrow search, the median ratio of CONGA-to-Percolator discoveries is consistently greater than 1 across FDR thresholds from 1% to 5%. Note that to ensure a fair comparison Percolator’s FDR analysis was also conducted with dynamic-level competition (see Percolator settings section).

Lastly, we compared the number of discoveries in the 24 peptide fractions from the HEK293 dataset reported by CONGA with those reported by TDC and Percolator (Supplementary Figure S5). We observe similar results to that of the PRIDE-20 results. Notably, using the median at the 1% FDR level, CONGA discovers 20.87% more discoveries than Percolator using a narrow-search mode and 2.79% more discoveries than Percolator using an open-search mode.

### CONGA accurately detects chimeric spectra

A chimeric spectrum is generated when the isolation following the MS1 scan fails to separate coeluting but distinct peptides. During the subsequent fragmentation step, ions are generated from multiple parent peptides, and these ions are simultaneously recorded in the following MS2 scan. Those chimeric spectra often pose a challenge for traditional “one-spectrum-one-peptide” tools that do not account for them ^19^.

CONGA’s design allows it to inherently process chimeras, identifying their multiple generating peptides, while still rigorously controlling the FDR in the overall reported list of detected peptides. To understand how this works, recall that CONGA considers multiple matches for each spectrum: one from the narrow search and up to *n*_*t*_ = 5 from the open search. The subsequent filtering process (see Section 2) guarantees that all remaining PSMs are made of peptides that essentially account for different peaks of the experimental spectrum. This requirement allows CONGA to distinguish between a truly co-generating peptide and a neighbor or “hitchhiking” peptide, where the latter offers a good match simply because its theoretical spectrum happens to share many peaks with the generating peptide.

The multiple PSMs are then assigned to different groups, and CONGA’s group-wise FDR control ensures that the overall FDR level is below the selected threshold. In particular, for non-chimeric spectra the secondary matches that get through the filtering will typically be low-scoring and therefore below the specific group threshold, whereas for some of the chimeric spectra multiple peptides can score high enough to be above the group threshold. We refer to peptides that CONGA identifies using a single spectrum as “co-peptides.” During group construction CONGA only considers the narrow and the top 2 open PSMs (this is a user-defined parameter for which we used the default of *R*_*i*_ = 2 throughout our analyses); thus, in principle, a chimeric spectrum that is detected by CONGA can contribute to the identification of up to three peptides in three (more generally *R*_*i*_ + 1) distinct groups, although in practice, using a 1% FDR threshold this seems to be an extremely rare event. Note that a PSM that comes up in the open search can still have an essentially 0 mass-modification, so CONGA can detect chimeric spectra regardless of whether each of the generating co-peptides includes a PTM.

To better understand CONGA’s ability to detect chimeric spectra, we set out to quantify the proportion of accepted peptides that are discovered via chimeric spectra. For this analysis, we consider the 24 spectrum files of the HEK293 dataset ^1^, and we focus on the MSFragger search results. We observe that at a 1% FDR threshold a median of 4.96% of the total peptides discovered by CONGA were identified thanks to chimeric spectra, and at 5% the median increases to 6.18% (Figure 4).

**Figure 4:**
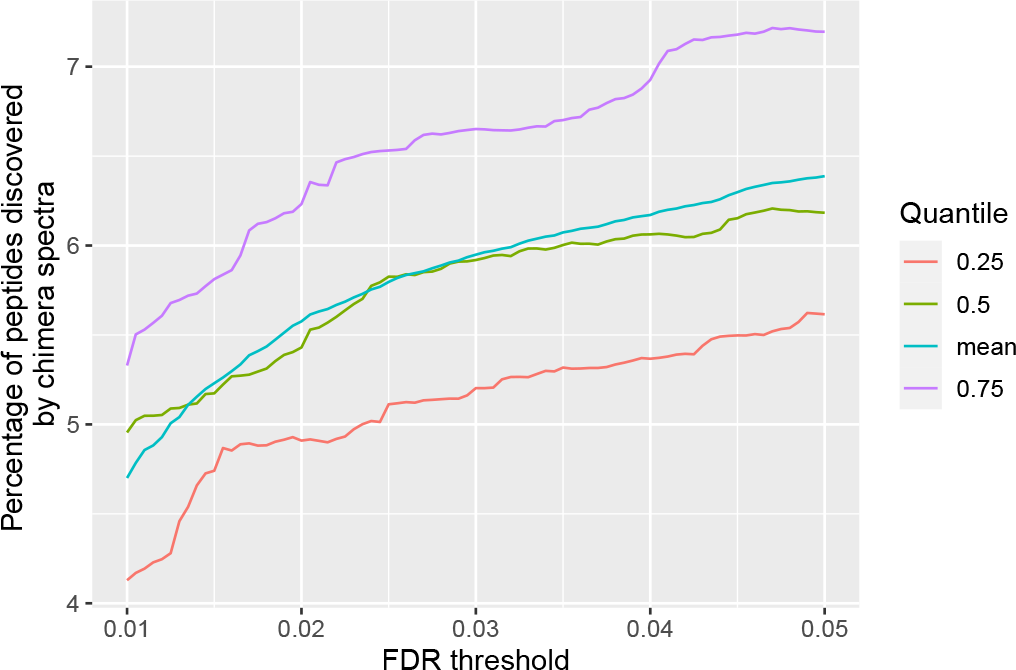
The percentage of peptides CONGA discovers through chimeric spectra. The quartiles and mean of the percentage of peptides detected by CONGA for which the top-scoring PSM is from a presumed chimeric spectra, plotted as a function of the FDR level. The quartiles and mean are taken over the 24 HEK293 spectrum files, and the percentages are taken with respect to the number of peptides discovered at the same FDR level by CONGA.

Although CONGA provides theoretical guarantees related to FDR control, we wanted to further validate the co-peptides in comparison to peptides detected using standard TDC applied to a narrow window search. To this end, we used two types of predictions from the machine learning tool Prosit ^21^: predicted peak intensities and predicted retention times. We hypothesized that the co-peptides should be indistinguishable from the peptides detected via TDC in terms of the accuracy of the predicted retention time and relative peak intensities. We tested this hypothesis on the representative HEK293 spectrum file (Section 3; HEK293 searches), collecting all of the co-peptides from CONGA and the peptides from MSFragger at a 1% FDR threshold, and using Prosit to predict peak intensities and retention times for both sets of peptides. Because Prosit generally does not allow for peptides with mass modifications as inputs, we only considered co-peptides where both have a mass that is no more than 1 Da away from that of the corresponding spectrum’s precursor.

We find that the accuracies of the Prosit predictions, for both peak intensities and retention times, are highly similar between the CONGA co-peptides and the MSFragger peptides (Figure 5). For the peak intensity prediction, visually there is very little that distinguishes between the two sets of discovered peptides (Figure 5A–B), and moreover, the overall correlation between the experimental and theoretical spectra in the two groups is essentially the same. For the retention time prediction, it is evident from Figure 5C–D that the relationship between the Prosit-predicted and experimental retention time is non-linear. Accordingly, we fitted a LOESS model, and we also used the Spearman correlation rather than the usual Pearson correlation. Again, we find the fit between the predicted and actual retention times seems to be invariant to whether the peptide was one identified by a presumed chimeric spectrum. The two corresponding Spearman correlation coefficients are also very close: 0.978 for the co-peptides and 0.972 for the TDC-discovered peptides. We note that the experimental retention time typically does not coincide with the apex retention time during peptide elution. Consequently, the Prosit predicted retention time may not agree exactly with the experimental one. Regardless, our aim is to only compare the fits of the co-peptides from CONGA and the narrow peptides from MSFragger.

**Figure 5:**
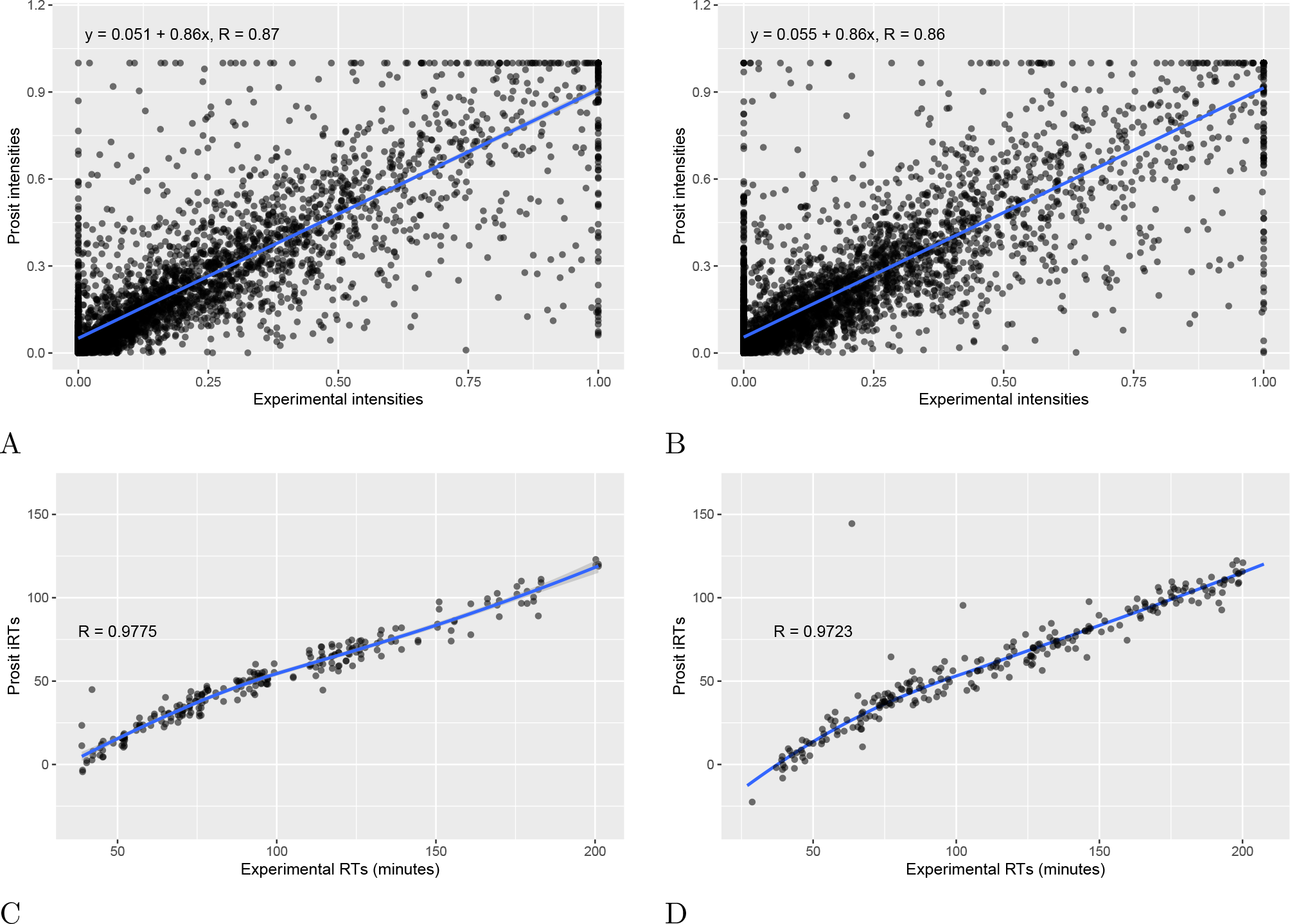
Comparisons of Prosit-predicted and observed peak intensities and retention times. (A) Scatter plot of experimental (x-axis) versus Prosit-predicted (y-axis) peak intensities from co-peptides discovered by CONGA at 1% FDR. (B) Similar scatter plot as panel A, but for peptides discovered by MSFragger in a narrow-window search at 1% FDR. For visualization, we only plot a randomly selected subset of the peak pairs, but the fitted line and correlation are computed over the entire set of peaks. (C) Scatter plot of observed versus Prosit-predicted retention times for co-peptides discovered by CONGA at 1% FDR. Note that technically we used Prosit’s normalized retention time indices (which can be negative), but for simplicity we refer to them as predicted retention times ^20^. (D) Similar plot as panel C, but for peptides discovered by MSFragger in a narrow-window search at 1% FDR. Again, panel D only includes a randomly selected subset of points, but the LOESS-fitted curve and the Spearman correlation are computed over the entire set of peptides in both C and D.

Finally, in Supplementary Figures S6 and S7 we used PDV ^22^ to manually annotate some of those presumed chimeric spectra identified by CONGA.

### CONGA uses pyAscore to help localize the mass modification

To illustrate CONGA’s use of pyAscore in localizing the modification we considered a representative HEK293 dataset that we randomly selected from the 24 HEK293 spectrum files (Section 3; HEK293 searches), and randomly further chose three PSMs in CONGA’s augmented list that scored above their group-threshold that coincide with acetylation, oxidation and phosphorylation (up to 0.05 Da). We then used PDV ^22^ to visually compare the pyAscore localizations reported in CONGA’s augmented list with the unmodified peptide from CONGA’s 1% FDR-controlled list (Supplementary Figure S1).

## 5. Discussion

Open search is an increasingly popular approach to detect peptides that harbor unexpected post-translational modifications. However, applying both open and narrow search to the same data, followed by TDC, often yields fewer discovered peptides for the open search than the narrow one. Only when applying a post-processor such as Percolator or PeptideProphet do we see a fairly consistent improvement using an open search. On the other hand, we demonstrated recently that both Percolator and PeptideProphet appear to underestimate the FDR in our entrapment experiments ^3^.

CONGA is designed as an alternative: it combines the results from the open and narrow searches in an approach that is built on a recently developed extension of the canonical TDC. CONGA’s control of the FDR draws on rigorous theoretical results; however, in practice, some of the necessary theoretical assumptions are only approximately satisfied. For example, the target database can often contain neighbors to some in-sample peptides, and such neighbors can compromise the assumption that an incorrect peptide is equally likely to be a target or a decoy. With its PSM-filtering as well as dynamic-level competition CONGA takes steps to mitigate the impact of neighbors.

Notably, in practice we see that CONGA apparently controls the FDR in the same entrapment experiments in which Percolator and PeptideProphet seemed to have failed ^3^. Interestingly, in spite of its tighter FDR control, CONGA’s power is comparable to, if not marginally better, than Percolator (we did not compare CONGA’s power with PeptideProphet’s). This is particularly surprising given that Percolator uses significantly more information than CONGA does, and at the same time it suggests a promising avenue for future gains by incorporating into CONGA the additional features that Percolator uses.

When introducing Group-walk we showed that we can use the mass difference between the precursor and the peptide to increase the power of the open search. However, the resulting open analysis occasionally fell significantly behind the results of narrow-TDC (Supplementary Figure S8). The current approach addressed that problem by co-opting the narrow search results into the analysis, allowing for multiple PSMs per spectrum, and applying a significantly more sophisticated, and automated, group construction procedure.

CONGA is a post-processor that provides rigorous FDR control by combining narrow and open searches. On its own, it does not try to localize the difference between the theoretical and observed mass of its reported discoveries. However, in principle, it can still be used to combine a narrow search with a localization-aware open search. We tested this capability with the recently introduced version of MSFragger, which matches the peaks of fragment ions containing up to one unknown PTM in the experimental spectrum to fragment ions in the theoretical peptide^23^. Notably, this approach has the potential to significantly improve the open search analysis, and therefore it is fair to ask whether it makes CONGA redundant. In Supplementary Figure S9 we demonstrate that even this localization-aware open search procedure benefits from using CONGA’s post-processing over TDC.

MetaMorpheus is another tool that attempts to localize modifications ^24^. Specifically, its enhanced G-PTM-D discovery component employs a two-search strategy, where the first utilizes a “multi-notch” search that considers a list of pre-determined possible mass modifications (unlike CONGA it cannot find unexpected mass modifications). It then uses a notch- or mass-specific FDR control to determine which of the observed modifications to use to modify the database so as to include all those possible modifications. Finally, it uses a second search, essentially an isotope error-tolerant narrow search of the spectra against the modified database. CONGA is not compatible with this two-stage procedure. Moreover, to the best of our understanding there is no theoretical foundation that supports the FDR control of the overall procedure, and some intermediate steps are particularly problematic. Specifically, we have previously showed that applying FDR analysis on each group does not control the FDR when the groups are combined; indeed, that was the motivation for our Group-walk procedure ^7^. Similarly, relying on a reversed, or for that matter any standardly generated decoy, cannot guarantee FDR control at the localization level. This is because of the same neighbors problem that we discussed in Section 2 and is the reason why CONGA employs dynamic-level competition.

CONGA requires as input a pairing between targets and decoys, which unfortunately is ill-defined if the decoy peptides are generated by first reversing or shuffling the entire protein. For this reason, CONGA only supports Tide, Comet and MSFragger, and the latter only if the target-decoy database is prepared beforehand, as explained in the last paragraph of Section 3 (PRIDE-20 searches).

We finish with a couple of tangential comments. The first is regarding the threshold (*τ* := 0.05) that we used for eliminating neighboring peptide matches in our initial filtering phase of CONGA. This threshold is likely lower than necessary for removing neighboring peptide matches. This low threshold incurs the cost of potentially throwing away some correctly matched co-peptides due to chimeric spectra. In fact, similar peptide sequences are more likely to co-elute, and so any reasonable amount of filtering will likely throw away such matches. However, setting it so low allows us to throw away many incorrect PSMs, and we found that the low threshold overall delivered more discoveries than the higher threshold of *τ* = 0.25 (data not shown). We note that detection of highly similar co-peptides with proper FDR control is a challenging task for CONGA, as well as for other post-processors, and is an interesting problem for future work.

As explained in Section 2, CONGA’s dynamic-level competition is motivated by theoretical FDR-control considerations. CONGA’s analysis switches between peptide-level or the representative-level depending on whether variable modifications are considered. However, Supplementary Figure S10A shows that in the 10 PRIDE-20 datasets whose analysis used variable modifications, there was little difference between the representative-level and peptide-level analyses. Specifically, we examined the power loss due to switching between the peptide and respresentative levels. Using CONGA at the 1% FDR level, we see median losses of 0.134% (Tailor) and 0.208% (XCorr), while at the 5% FDR the corresponding median percentage losses are 0.450% and 0.543%. Panel B of the same figure shows that the effect is even milder when narrow TDC is used instead. This is reassuring given that all results comparing CONGA to *all other methods* were done at the representative-level when variable modifications were used.

## SUPPORTING INFORMATION

The following supporting information is available free of charge at ACS website https://pubs.acs.org/.

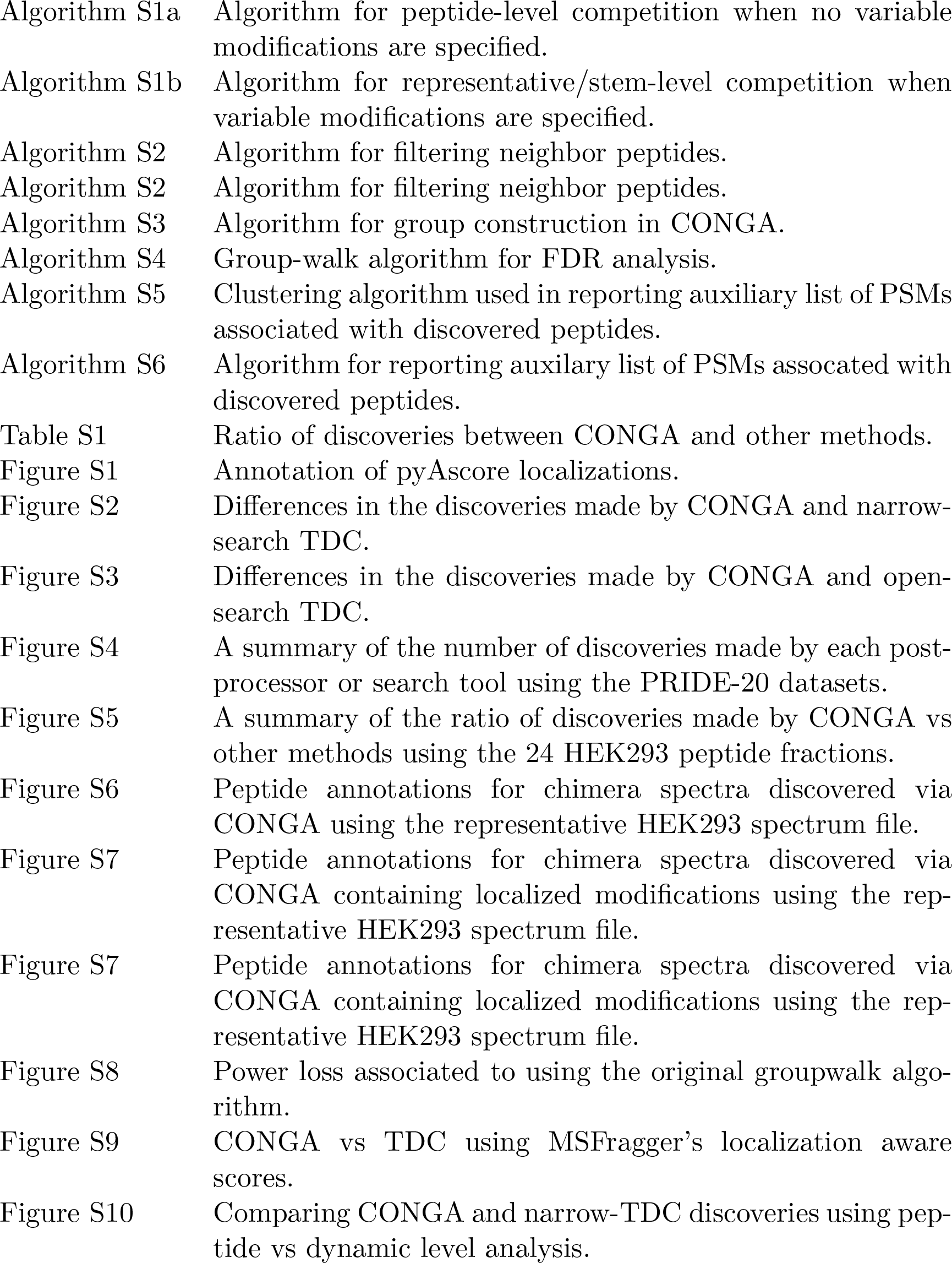

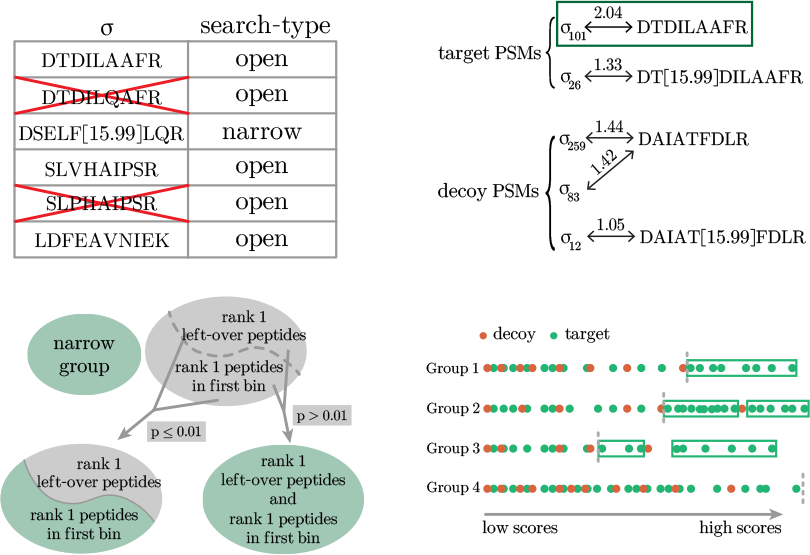

For TOC Only

## Notes

https://github.com/freejstone/CONGA

### Competing Interest Statement

The authors have declared no competing interest.

### Summary of Updates

This version represents a significant update to the content of the paper. The paper is now organized differently with a new section called Approach providing a high-level description of CONGA. A number of new analyses were added and the exposition was improved throughout the paper.

