## Supplementary Information for "Analysis of tandem mass spectrometry data with CONGA: Combining Open and Narrow searches with Group-wise Analysis"

### Supporting Information

| Table of Contents | |
|---|---|
| Algorithm S1a | Algorithm for peptide-level competition when no variable modifications are specified. |
| Algorithm S1b | Algorithm for representative/stem-level competition when variable modifications are specified. |
| Algorithm S2 | Algorithm for filtering neighbor peptides. |
| Algorithm S2 | Algorithm for filtering neighbor peptides. |
| Algorithm S3 | Algorithm for group construction in CONGA. |
| Algorithm S4 | Group-walk algorithm for FDR analysis. |
| Algorithm S5 | Clustering algorithm used in reporting auxiliary list of PSMs associated with discovered peptides. |
| Algorithm S6 | Algorithm for reporting auxilary list of PSMs assocated with discovered peptides. |
| Table S1 | Ratio of discoveries between CONGA and other methods. |
| Figure S1 | Annotation of pyAscore localizations. |
| Figure S2 | Differences in the discoveries made by CONGA and narrow-search TDC. |
| Figure S3 | Differences in the discoveries made by CONGA and open-search TDC. |
| Figure S4 | A summary of the number of discoveries made by each post-processor or search tool using the PRIDE-20 datasets. |
| Figure S5 | A summary of the ratio of discoveries made by CONGA vs other methods using the 24 HEK293 peptide fractions. |
| Figure S6 | Peptide annotations for chimera spectra discovered via CONGA using the representative HEK293 spectrum file. |
| Figure S7 | Peptide annotations for chimera spectra discovered via CONGA containing localized modifications using the representative HEK293 spectrum file. |
| Figure S7 | Peptide annotations for chimera spectra discovered via CONGA containing localized modifications using the representative HEK293 spectrum file. |
| Figure S8 | Power loss associated to using the original groupwalk algorithm. |
| Figure S9 | CONGA vs TDC using MSFragger's localization aware scores. |
| Figure S10 | Comparing CONGA and narrow-TDC discoveries using peptide vs dynamic level analysis. |

**Algorithm S1a PSM-and-peptide** from [1]

**Input:**
- $(\mathcal{T}, \mathcal{D})$ - the target and decoy databases;
- $\{(\sigma, \pi_T^\sigma, \pi_D^\sigma, S_T^\sigma, S_D^\sigma) : \sigma \in \Sigma\}$ - the set of experimental spectra, their target and decoy matches, and their associated scores;
- `TD_pairs` - a mapping that pairs every target peptide to its decoy;
- $\alpha$ - the FDR threshold;

**Output:** $R$ - a discovery list containing a subset of the target peptides and their scores

**for** $\sigma \in \Sigma$ **do** ▷ PSM-level competition
  **if** $S_T^\sigma > S_D^\sigma$ **then** ▷ if $S_T^\sigma = S_D^\sigma$, randomly break ties
    $(\pi^\sigma, S^\sigma) \leftarrow (\pi_T^\sigma, S_T^\sigma)$
  **else**
    $(\pi^\sigma, S^\sigma) \leftarrow (\pi_D^\sigma, S_D^\sigma)$
  **end if**
**end for**
**for** $\pi \in \mathcal{T} \cup \mathcal{D}$ **do** ▷ obtaining the best score associated with each peptide $\pi$
  $S_\pi \leftarrow \max\{S^\sigma \mid \pi^\sigma = \pi\}$ ▷ where $\max(\emptyset) = -\infty$
**end for**
$WP \leftarrow \emptyset$ ▷ triplets characterizing the winning target/decoy peptides $\pi$
**for** $\pi \in \mathcal{T}$ **do** ▷ peptide-level competition
  **if** $S_\pi > S_{\texttt{TD\_pairs}(\pi)}$ **then** ▷ if $S_\pi = S_{\texttt{TD\_pairs}(\pi)}$, randomly break ties
    $WP \leftarrow WP \cup (\pi, S_\pi, 1)$ ▷ add a target winning peptide
  **else**
    $WP \leftarrow WP \cup (\texttt{TD\_pairs}(\pi), S_{\texttt{TD\_pairs}(\pi)}, -1)$ ▷ add a decoy winning peptide
  **end if**
**end for**
Order the triplets of $WP = \{(\pi_i, W_i, L_i)\}$ so that $W_1 \geq \cdots \geq W_n$ ▷ ties are randomly broken
$K \leftarrow \max\{k : (\#\{i \leq k : L_i = -1\} + 1)/(\#\{i \leq k : L_i = 1\} \vee 1) \leq \alpha\}$ ▷ FDR analysis
**return** $R \leftarrow \{(\pi_i, W_i) \mid i \leq K, L_i = 1\}$

**Algorithm S1b PSM-and-peptide adapted for variable modifications**

**Input:**
- $(\mathcal{T}^s, \mathcal{D}^s)$ - the target and decoy databases containing only the **stem** form peptides;
- $\{(\sigma, \pi_T^\sigma, \pi_D^\sigma, S_T^\sigma, S_D^\sigma) : \sigma \in \Sigma\}$ - the set of experimental spectra, their target and decoy matches, and their associated scores;
- `TD_pairs` - a mapping that pairs every target stem peptide to its decoy;
- `remove_mods` - a mapping that takes a peptide and returns its stem form;
- $\alpha$ - the FDR threshold;

**Output:** $R$ - a discovery list containing a subset of the (representative) target peptides and their scores

**for** $\sigma \in \Sigma$ **do** ▷ PSM-level competition
  **if** $S_T^\sigma > S_D^\sigma$ **then** ▷ if $S_T^\sigma = S_D^\sigma$, randomly break ties
    $(\pi^\sigma, S^\sigma, \mathring{\pi}^\sigma) \leftarrow (\pi_T^\sigma, S_T^\sigma, \texttt{remove\_mods}(\pi_T^\sigma))$
  **else**
    $(\pi^\sigma, S^\sigma, \mathring{\pi}^\sigma) \leftarrow (\pi_D^\sigma, S_D^\sigma, \texttt{remove\_mods}(\pi_D^\sigma))$
  **end if**
**end for**
**for** $\mathring{\pi} \in \mathcal{T}^s \cup \mathcal{D}^s$ **do** ▷ obtaining the best score associated with each stem peptide $\mathring{\pi}$
  $S_{\mathring{\pi}} \leftarrow \max\{S^\sigma \mid \mathring{\pi}^\sigma = \mathring{\pi}\}$ ▷ where $\max(\emptyset) = -\infty$
  $\pi_{\mathring{\pi}} \leftarrow$ randomly select from $\{\pi^\sigma \mid S^\sigma = S_{\mathring{\pi}}, \mathring{\pi}^\sigma = \mathring{\pi}\}$ ▷ NULL if $S_{\mathring{\pi}} = -\infty$
**end for**
$WS \leftarrow \emptyset$ ▷ triplets characterizing the winning target/decoy representatives
**for** $\mathring{\pi} \in \mathcal{T}^s$ **do** ▷ representative-level competition
  **if** $S_{\mathring{\pi}} > S_{\texttt{TD\_pairs}(\mathring{\pi})}$ **then** ▷ if $S_{\mathring{\pi}} = S_{\texttt{TD\_pairs}(\mathring{\pi})}$, randomly break ties
    $WS \leftarrow WS \cup (\pi_{\mathring{\pi}}, S_{\mathring{\pi}}, 1)$ ▷ add a target winning representative
  **else**
    $WS \leftarrow WS \cup (\texttt{TD\_pairs}(\pi_{\mathring{\pi}}), S_{\texttt{TD\_pairs}(\mathring{\pi})}, -1)$ ▷ add a decoy winning representative
  **end if**
**end for**
Order the triplets of $WS = \{\pi_i, W_i, L_i)\}$ so that $W_1 \geq \cdots \geq W_n$ ▷ ties are randomly broken
$K \leftarrow \max\{k : (\#\{i \leq k : L_i = -1\} + 1)/(\#\{i \leq k : L_i = 1\} \vee 1) \leq \alpha\}$ ▷ FDR analysis
**return** $R \leftarrow \{(\pi_i, W_i) \mid i \leq K, L_i = 1\}$

**Algorithm S2** Filter neighboring peptides

**Input:** • $\{(\sigma, \pi_j^\sigma) : j = 1, \ldots, k^\sigma, \sigma \in \Sigma\}$ - the collection of PSMs represented as pairs of experimental spectra and matching peptides;
• $\tau$ - a similarity threshold (we used $\tau := 0.05$)

**Output:** $\{(\sigma, \pi_j^\sigma) : j \in J^\sigma, \sigma \in \Sigma\}$ - a neighbor-filtered subset of the PSMs

```
for σ ∈ Σ do
    i ← 1
    J^σ ← {1}        ▷ J^σ contains the indices of retained peptides associated with σ
    while i + 1 ≤ k^σ do
        Using equation (1) compute the similarities {t_j^σ : j ∈ J^σ} between the peptide π_{i+1}^σ and
            each of the peptides in {π_j^σ : j ∈ J^σ}
        i ← i + 1
        if t_j^σ < τ for all j ∈ J^σ then
            J^σ ← J^σ ∪ {i + 1}
        end if
    end while
end for
return {(σ, π_j^σ) for j ∈ J^σ, for σ ∈ Σ}
```

**Algorithm S3** Group construction

**Input:**
- $\{(W_i, N_i, R_i, \delta_i) : i = 1, \ldots, m\}$ - tuples characterizing the winning peptides (winning score, narrow label, PSM rank, mass difference);
- $K$ - a window size used by Group-walk (usually $K = 40$);
- $\lambda$ a precursor bin width (usually $\lambda = 1.0005079/4$);
- $\beta \in (0, 1)$ - a threshold value used for merging groups (usually $\beta = 0.01$)
- $k$ - the number of top PSMs in the open-search file considered by each spectrum

**Output:** $G_i$ - the group labels for all winning peptides

Partition the peptides with $N_i = 0$ into bins according to the mass differences $\delta_i$ using bins of size $\lambda$ centred at multiples of $\lambda$.
Order the bins according to the number of peptides with $R_i = 1$ within each bin from largest to smallest.
$b_i \leftarrow$ the rank of the bin containing the $i$th peptide for $i = 1, \ldots, m$.
$b \leftarrow \max\{b_i\}$ ▷ largest ranked bin for top 1 open-search peptides
$G_i \leftarrow 0$ for $i = 1, \ldots, m$ such that $N_i = 1$. ▷ create the group-labels for narrow-search peptides
$G_i \leftarrow 1$ for $i = 1, \ldots, m$ such that $N_i = 0$. ▷ initalize the group-labels for open-search peptides
$n_g \leftarrow 1$ ▷ initialize the number of groups for open-search peptides
$i \leftarrow 1$
**while** $i \leq b$ **do**
  $C \leftarrow \{l \mid b_l = i, N_l = 0, R_l = 1\}$ ▷ defining the candidate set $C$ as a potential group
  **while** $|C| < 2K$ and $i < b$ **do**
    $i \leftarrow i + 1$
    $C \leftarrow C \cup \{l \mid b_l = i, N_l = 0, R_l = 1\}$
  **end while**
  **if** $i = b$ and $|C| < 2K$ **then**
    $G_l \leftarrow n_g$ for all $l \in C$ ▷ join to the previously defined group
    break
  **else if** $i = b$ and $|C| \geq 2K$ **then**
    $n_g \leftarrow n_g + 1$
    $G_l \leftarrow n_g$ for all $l \in C$ ▷ create new group
    break
  **end if**
  $L \leftarrow \{l \mid b_l > i, N_l = 0, R_l = 1\}$ ▷ the remaining set of top 1 peptides yet to be grouped
  $p \leftarrow$ `KS.TEST`$(\{W_l \mid l \in C\}, \{W_l \mid l \in L\})$ ▷ compare the scores between $C$ and $L$

*continues onto next page...*

**Algorithm S3** Group construction (continued)

...

**if** $p \leq \beta$ **then**
  **if** $n_g = 1$ **then**
    $n_g \leftarrow n_g + 1$
    $G_l \leftarrow n_g$ for all $l \in C$
  **else**
    Let $\Pi : \{2, \ldots, n_g\} \to \{2, \ldots, n_g\}$ be a random permutation
    $p_l \leftarrow 0$ for $l = 2, \ldots, n_g$
    **for** $j \in \{2, \ldots, n_g\}$ **do**
      $T \leftarrow \{l \mid G_l = \Pi(j)\}$
      $p_j \leftarrow \texttt{KS.TEST}(\{W_l \mid l \in C\}, \{W_l \mid l \in T\})$
      **if** $p_j > \beta$ **then**
        $G_l \leftarrow \Pi(j)$ for $l \in C$ ▷ join $C$ to an existing group if they are similar
        break
      **end if**
    **end for**
    **if** $p_l \leq \beta$ for all $l \in \{2, \ldots, n_g\}$ **then**
      $n_g \leftarrow n_g + 1$
      $G_l \leftarrow n_g$ for all $l \in C$ ▷ $C$ defines a new group if different to all other groups
    **end if**
  **end if**
**else**
  $n_g \leftarrow n_g + 1$
  $G_l \leftarrow n_g$ for all $l \in C \cup L$ ▷ combine $C$ and $L$ into a group if they are similar
  break
**end if**
**end while**
$H \leftarrow \{i \mid b_i \leq n_g, N_i = 0, R_i > 1\}$ ▷ peptides ranked from 2 to $k$ that belong to a bin that is responsible for creating a group
$J \leftarrow \{i \mid b_i > n_g, N_i = 0, R_i > 1\}$ ▷ remaining peptides
**if** $|H| \geq 2K$ **then**
  $n_g \leftarrow n_g + 1$
  $G_i \leftarrow n_g$ for all $i \in H$ ▷ create a new group out of $H$
**else**
  $G_i \leftarrow n_g$ for all $i \in H$ ▷ join $H$ to the last group
**end if**
$G_i \leftarrow \texttt{NULL}$ for all $i \in J$
**return** $G_i$ for $i = 1, \ldots, m$

**Algorithm S4** Group-walk

**Input:** • $\alpha$ - the FDR threshold $\in (0,1)$;
• $K$ - the "look-back" window size;
• $\{L_i^g : g = 1, \dots, n_g, i = 1, \dots, m_g\}$ - $\pm 1$ target/decoy win labels corresponding to the *increasingly sorted* $m_g$ winning scores $W_i^g$ within each of the $n_g$ groups: $W_1^g \leq W_2^g \leq \dots \leq W_{m_g}^g$;

**Output:** $\bigcup_g \{i \geq k_g \mid L_i^g = 1\}$ - the discovery list

$k_g \leftarrow 1$ for $g \in \{1, \dots, n_g\}$ ▷ initialise the frontier vector
**while** there exists $g \in \{1, \dots, n_g\}$ such that $k_g \leq m_g$ **do**
    $D_g(k_g) = \sum_{i=k_g}^{m_g} 1_{\{L_i^g = -1\}}$ ▷ the # decoys above the frontier vector in each group
    $R_g(k_g) = \sum_{i=k_g}^{m_g} 1_{\{L_i^g = 1\}}$ ▷ the # targets above the frontier vector in each group
    $F\hat{D}R \leftarrow \frac{1 + \sum_g D_g(k_g)}{1 \vee \sum_g R_g(k_g)}$ ▷ the estimated false discovery proportion
    **if** $F\hat{D}R \leq \alpha$ **then**
        break
    **else**
        **if** there exists $g \in \{1, \dots, n_g\}$ such that $k_g \leq K$ **then**
            $l \leftarrow$ randomly select from $\{l \mid k_l = \min_g k_g\}$
            $k_l \leftarrow k_l + 1$
        **else**
            **for** $g \in \{1, \dots, n_g\}$ **do**
                $w_g \leftarrow D_g(k_g - K) - D_g(k_g)$ ▷ the number of decoys in the last $K$ labels
            **end for**
            $l \leftarrow$ randomly select from $\{l \mid w_l = \max_g w_g\}$
            $k_l \leftarrow k_l + 1$ ▷ increase the frontier according to $w_g$
        **end if**
    **end if**
**end while**
**return** $\bigcup_g \{i \geq k_g \mid L_i^g = 1\}$

**Algorithm S5** Precursor mass clustering (`precursor_clustering`)

**Input:** • $\{(\rho_i, c_i) : i = 1, \dots, n\}$ - a set of precursor masses and charge states associated with a given stem;
• $(w_l, w_u)$ - the experimental isolation window given as a pair of values;

**Output:** $\{P_j\}$ - the clustered precursors;

Reorder the precursors so that $\rho_1 \leq \cdots \leq \rho_n$
$j \leftarrow 1$
$P_j \leftarrow \{1\}$
**for** $i \in \{2, \dots, n\}$ **do**
    **if** $\rho_i \leq \rho_{i-1} + \max\{c_{i-1} w_u, c_i w_l\}$ **then**
        $P_j \leftarrow P_j \cup \{i\}$
    **else**
        $j \leftarrow j + 1$
        $P_j \leftarrow \{i\}$
    **end if**
**end for**
**return** $\{P_j\}$

**Algorithm S6** Generating the augmented list of target PSMs with variants of the discovered peptides

**Input:**
- $\{\pi_i^* : i = 1, \ldots, M\}$ - the set of peptide discoveries in stem (unmodified) form;
- $\{(\sigma_i, \pi_i, mod_i, \rho_i, c_i, S_i, N_i, R_i, \delta_i) : i = 1, \ldots, m\}$ - a set of tuples characterizing the $m$ target PSMs that survived the neighbor filtering and whose rank is $\leq 2$, where $\sigma_i$ =spectrum, $\pi_i$ =stem (unmodified) peptide, $mod_i$ =modification, $\rho_i$ =precursor mass, $c_i$ =precursor charge, $S_i$ =PSM score, $N_i$ =narrow label, $R_i$ =PSM rank, and $\delta_i$ =mass-difference;
- $(w_l, w_u)$ - the experimental isolation window given as a pair of values;
- `get_groups` - the mapping which associates the narrow label, PSM rank, and delta mass $(N, R, \delta)$ to one of $n_g$ group labels $g_1, \ldots, g_{n_g}$ according to the group construction step (Algorithm S3);
- $\{T_{g_i} : i = 1, \ldots, n_g\}$ - the group score thresholds, one for each of the $n_g$ groups;

**Output:**
- $J$ - the augmented list of PSMs to be reported;
- $\{a_i : i \in J\}$ - labels indicating if the corresponding PSM is greater than or equal to its associated group score threshold ($a_i = 1$) or if it is below ($a_i = 0$);

$a_i \leftarrow 0$ for all $i = 1, \ldots, m$
$J \leftarrow \emptyset$ ▷ $J$ is the indices of the PSMs to be returned
**for** $\pi^* \in \{\pi_1^*, \ldots, \pi_M^*\}$ **do**
    $I \leftarrow \{i \mid \pi^* = \pi_i\}$
    $mods \leftarrow \{mod_i \mid i \in I\}$
    **for** $i \in I$ **do**
        $G_i \leftarrow$ `get_groups`$(N = N_i, R = R_i, \delta = \delta_i)$
    **end for**
    $Cs \leftarrow$ `precursor_clustering`$(\{(\rho_i, c_i)\}_{i \in I}, (w_l, w_u))$ ▷ applying Algorithm S5
    **for** $C \in Cs$ and $mod \in mods$ **do**
        $S_x \leftarrow \max\{S_j \mid j \in C, mod_j = mod\}$
        **if** $S_x > -\infty$ **then** ▷ $\max\{\emptyset\} = -\infty$
            $l \leftarrow$ randomly select from $\{i \mid S_i = S_x, mod_i = mod\}$
            $J \leftarrow J \cup \{l\}$
            **if** $S_l \geq T_{G_l}$ **then**
                $a_l \leftarrow 1$
            **end if**
        **end if**
    **end for**
**end for**
**return** $\{(\sigma_i, \pi_i, mod_i, \rho_i, c_i, S_i, N_i, R_i, \delta_i, a_i)\}_{i \in J}$

| | Tide | | Comet | | MSFragger | | Percolator Tide | |
|---|---|---|---|---|---|---|---|---|
| Project ID | Narrow | Open | Narrow | Open | Narrow | Open | Narrow | Open |
| PXD008920 | 1.059 | 1.210 | 1.171 | 1.199 | 1.102 | 1.203 | 1.027 | 1.048 |
| PXD008996 | 1.055 | 1.261 | 1.262 | 1.316 | 1.084 | 1.329 | 1.020 | 1.071 |
| PXD010504 | 1.239 | 1.321 | 1.268 | 1.405 | 1.117 | 1.614 | 1.168 | 1.074 |
| PXD014277 | 1.055 | 1.093 | 1.064 | 1.106 | 1.057 | 1.107 | 1.009 | 1.017 |
| PXD016724 | 1.545 | 1.158 | 1.693 | 1.145 | 1.193 | 1.280 | 1.493 | 1.045 |
| PXD019354 | 1.177 | 1.218 | 1.381 | 1.244 | 1.191 | 1.283 | 1.137 | 1.043 |
| PXD024284 | 1.028 | 1.141 | 1.024 | 1.251 | 1.063 | 1.172 | 1.006 | 1.040 |
| PXD025130 | 1.122 | 1.249 | 1.112 | 1.155 | 1.157 | 1.256 | 1.046 | 1.021 |
| PXD029319 | 1.063 | 1.071 | 2.392 | 1.003 | 1.207 | 1.087 | 1.06 | 1.054 |
| PXD030118 | 1.155 | 1.143 | 1.134 | 1.143 | 1.169 | 1.177 | 1.110 | 1.046 |
| PXD002470 | 0.993 | 1.225 | 1.543 | 1.037 | 0.967 | 0.922 | 0.875 | 0.881 |
| PXD006856 | 1.694 | 1.069 | 1.563 | 1.367 | 1.578 | 1.204 | 1.671 | 1.021 |
| PXD010413 | 1.059 | 1.10 | 1.098 | 0.975 | 1.045 | 1.153 | 0.990 | 0.935 |
| PXD012528 | 1.122 | 1.313 | 1.168 | 1.977 | 1.098 | 1.274 | 1.161 | 0.974 |
| PXD012611 | 1.011 | 1.048 | 0.991 | 1.082 | 1.033 | 1.127 | 1.008 | 0.983 |
| PXD013274 | 1.029 | 1.063 | 1.029 | 1.059 | 1.087 | 1.095 | 1.027 | 1.031 |
| PXD019186 | 2.884 | 1.681 | 2.780 | 1.893 | 1.155 | 1.538 | 2.501 | 1.263 |
| PXD022257 | 1.259 | 1.111 | 1.510 | 1.121 | 1.177 | 1.070 | 1.234 | 0.998 |
| PXD023571 | 0.987 | 1.973 | 1.016 | 1.755 | 1.004 | 5.057 | 0.707 | 0.875 |
| PXD026895 | 1.247 | 1.264 | 1.258 | 1.218 | 1.310 | 1.318 | 1.188 | 1.058 |

Table S1: **CONGA vs other methods** The mean ratio of CONGA-to-other methods. For Tide-results, the mean ratio is taken over each dataset's 20 decoys, while Comet's and MSFragger's searches used a single reversed set of decoys. Notably the number of open search discoveries with MSFragger using PXD023571 was low (we believe due to the higher fragment tolerance required for this dataset).

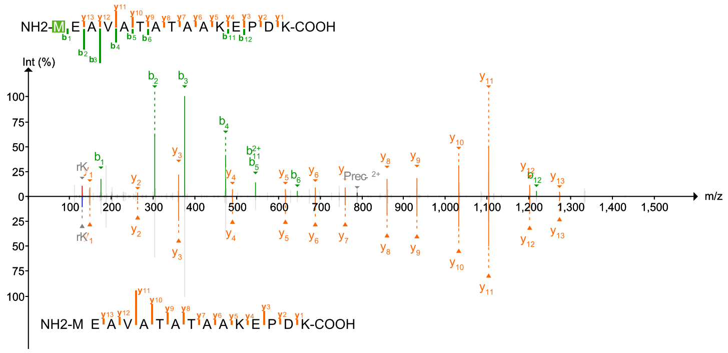


Scan 37354

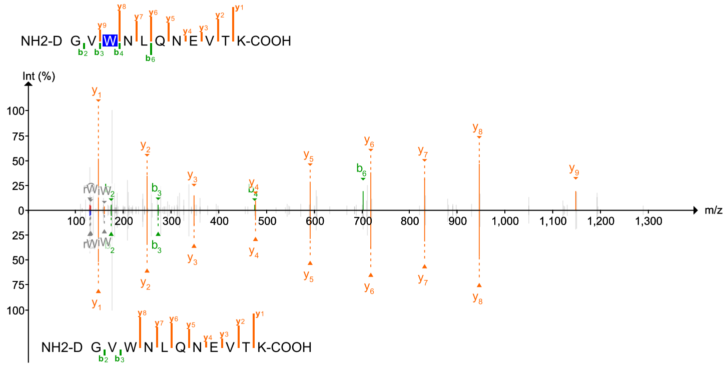


Scan 43833

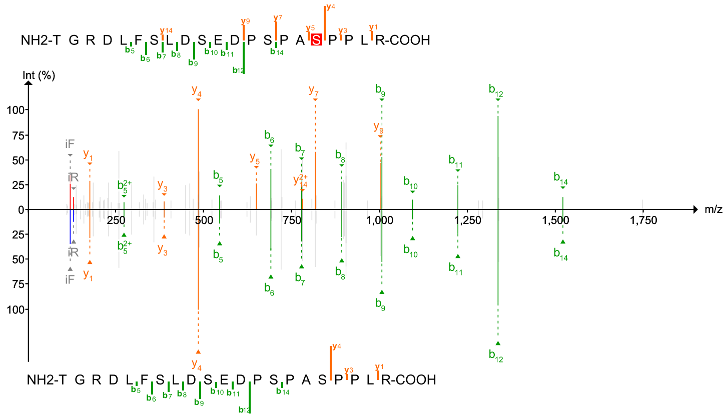


Scan 59837

Figure S1: **Peptide annotations for pyAscore localizations.** Mirror plots of three experimental spectra, annotated by the modified peptide localized using pyAscore, as reported in CONGA's augmented list (top), and the original unmodified peptide appearing in CONGA's 1% FDR-controlled list (bottom). The modifications detected are acetylation, oxidation and phosphorylation respectively. The scan numbers relate to the representative HEK293 dataset.

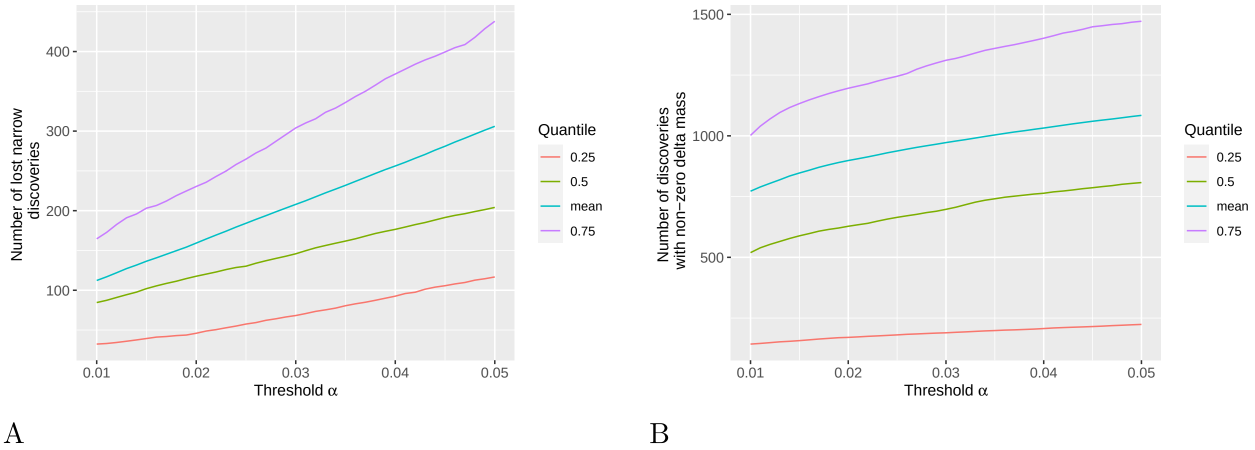


Figure S2: **Differences in CONGA discoveries and narrow TDC.** (A) The quartiles and mean of the average number of peptides discovered by narrow-TDC (PSM-and-peptide applied to Tide-generated narrow-search PSMs) that were lost in CONGA (applied to the open and narrow Tide generated PSMs). (B) The quartiles and mean of the average number of peptides discovered by CONGA outside of the narrow-window precursor-tolerance. The quartiles and mean are taken of the 20 PRIDE-20 datasets and the average percentage of lost peptides is taken over the 20 decoys generated per dataset.

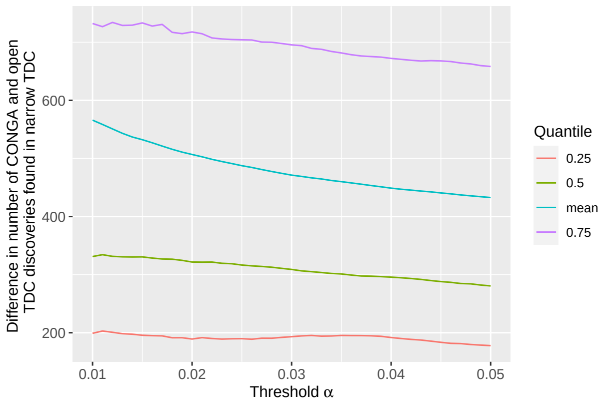


A

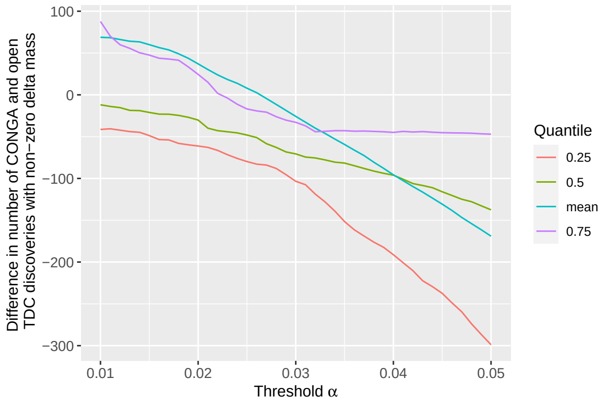


B

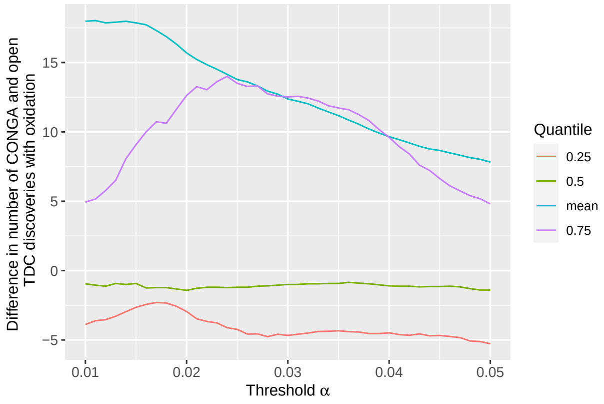


C

Figure S3: **Differences in CONGA discoveries and open TDC.** All plots were determined from Tide-search results. (A) The quartiles and mean of the average number of peptides simultaneously discovered by CONGA and narrow-TDC (PSM-and-peptide applied to Tide-generated narrow-search PSMs) minus the discoveries made by open- and narrow-TDC. (B) The quartiles and mean of the average difference in peptide discoveries between CONGA and open-TDC outside of the narrow-window precursor tolerance. (C) The quartiles and mean of the average difference in peptide discoveries between CONGA and open-TDC with delta-mass at 15.995 Da (± 0.2 Da). In all cases the quartiles and mean are taken of the 20 PRIDE-20 datasets and the averages are taken over the 20 decoys generated per dataset.

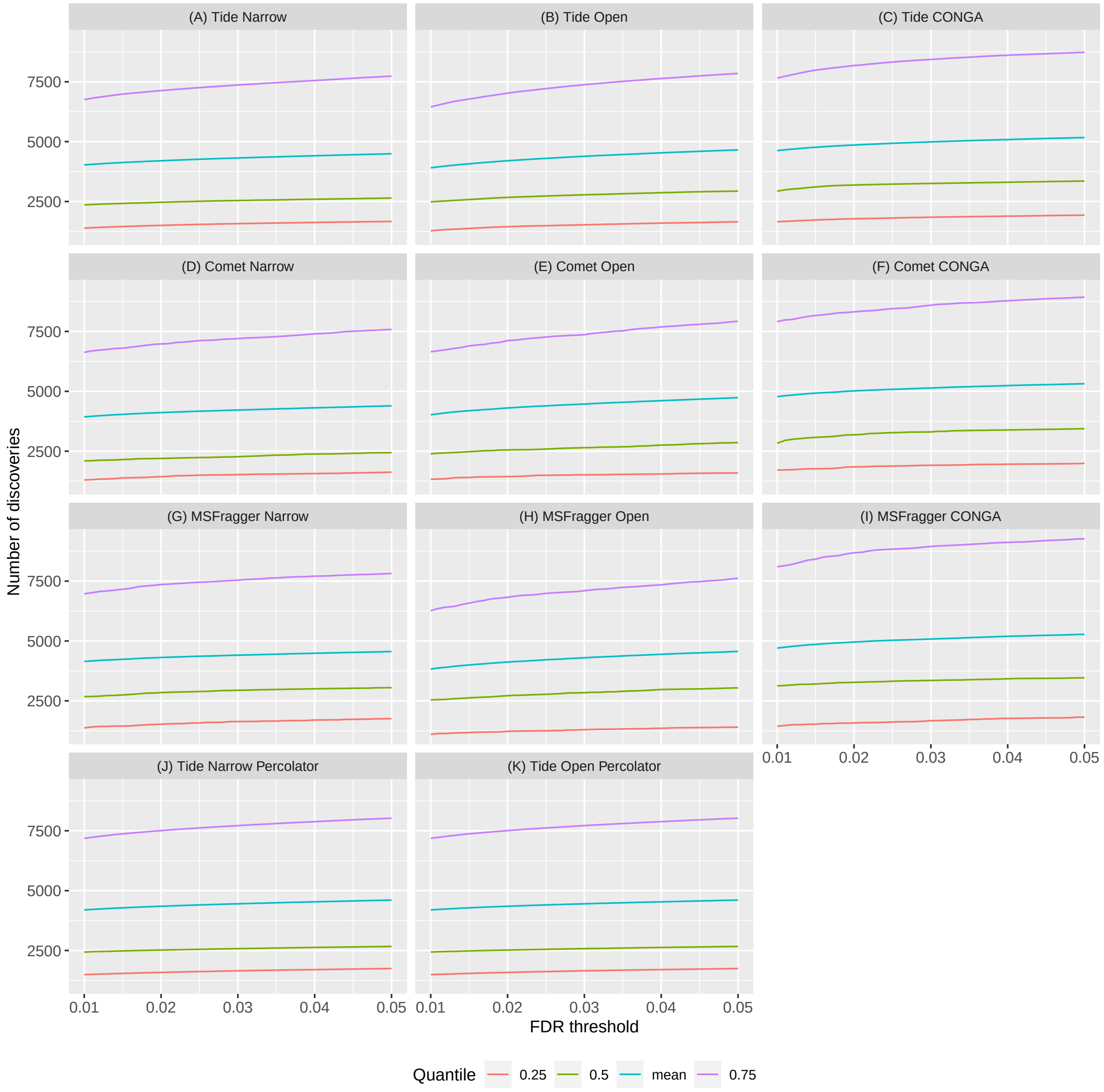


Figure S4: **The quartiles of the number of discoveries made by each search tool or post-processor.** The quartiles and mean, taken over the average number of discoveries using the 20 PRIDE-20 datasets. Each of the first three rows represents a different search engine (Tide, Comet, MSFragger), and the last row corresponds to Percolator applied to Tide-search results. The first column refer to narrow searches, the second column to open searches, and the last column to CONGA, combining the two search modes. For the results that use Tide, the average number of discoveries is taken with respect to each dataset's 20 decoys, while Comet's and MSFragger's searches use a single reversed set of decoys.

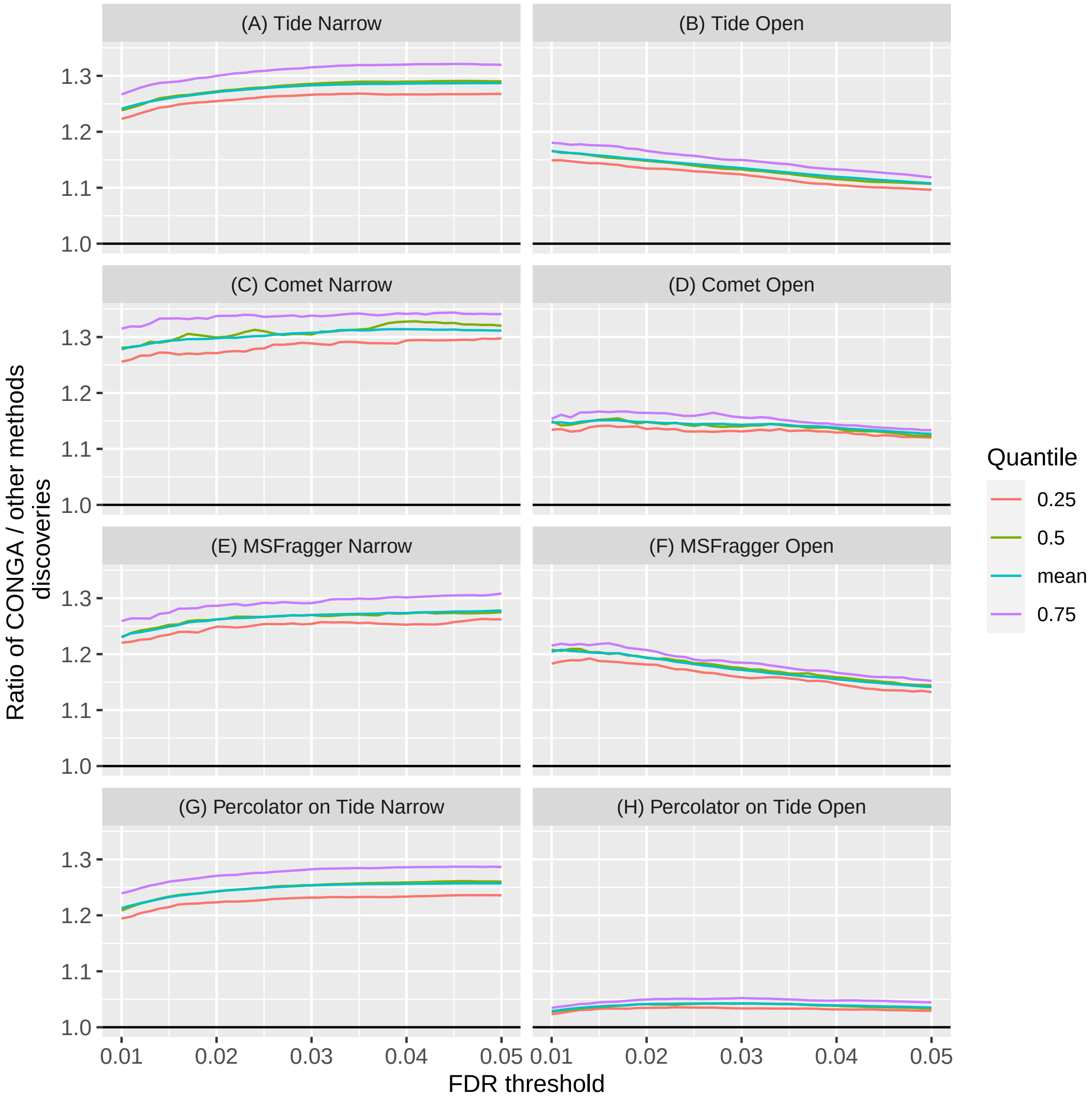


Figure S5: **CONGA detects more peptides than narrow or open searches applied to HEK293 data set.** The quartiles and mean, taken over the 24 HEK293 spectrum files, of the average ratio of CONGA-to-TDC discovered peptides. TDC was applied to (A) Tide search results in narrow mode, (B) Tide search results in open mode, (C) Comet search results in narrow mode, (D) Comet search results in open mode, (E) MSFragger search results in narrow mode and (F) MSFragger search results in open mode. Panels (G) and (H) are similar, but compare CONGA with Percolator applied to the Tide narrow and open search results, respectively. For results that use Tide the average ratio is taken with respect to each dataset's 20 decoys, while Comet's and MSFragger's searches used a single reversed set of decoys.

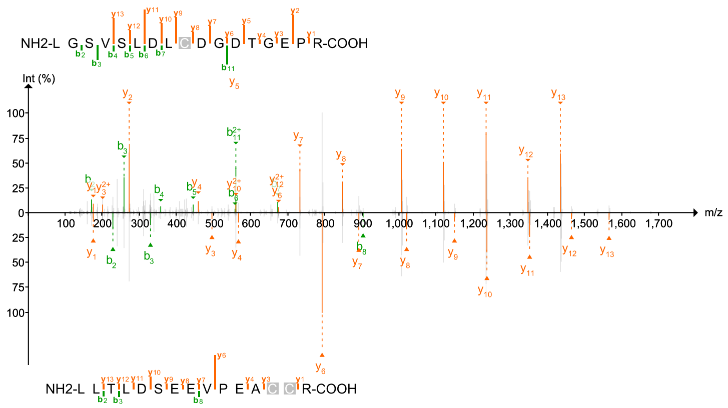


Scan 45852

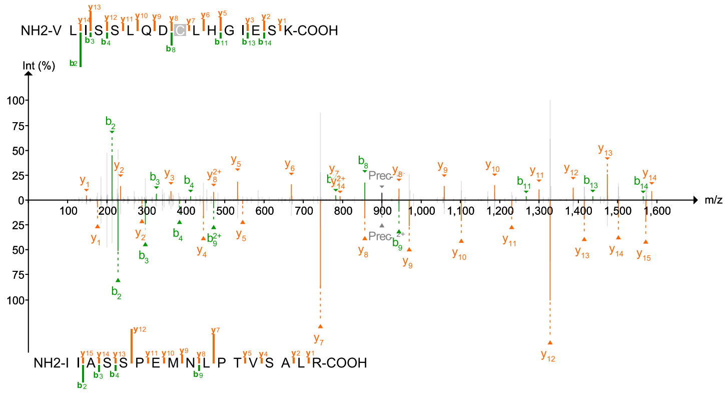


Scan 63526

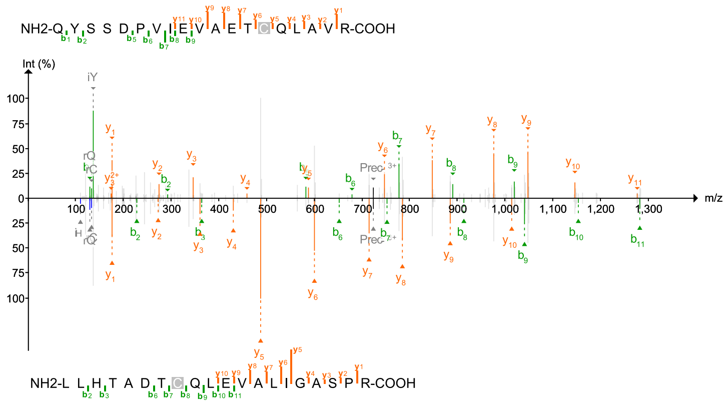


Scan 66043

Figure S6: **Peptide annotations for chimera spectra.** Mirror plots of three experimental spectra, annotated by the corresponding pairs of peptides discovered using CONGA at the 1% FDR level. The same experimental spectrum is reflected horizontally, with matched b-ions annotated in green and matched y-ions annotated in orange. The scan numbers relate to the representative HEK293 dataset.

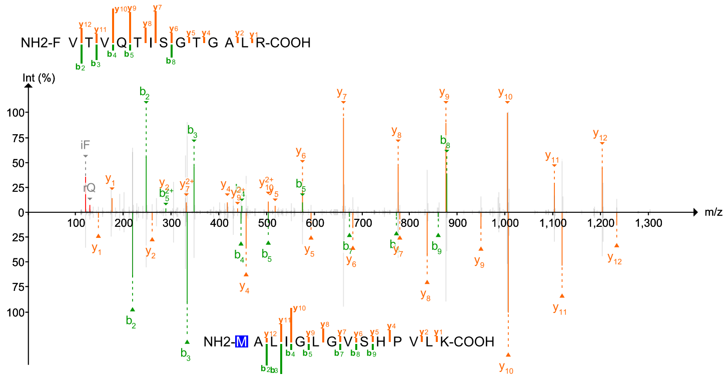


Scan 47880

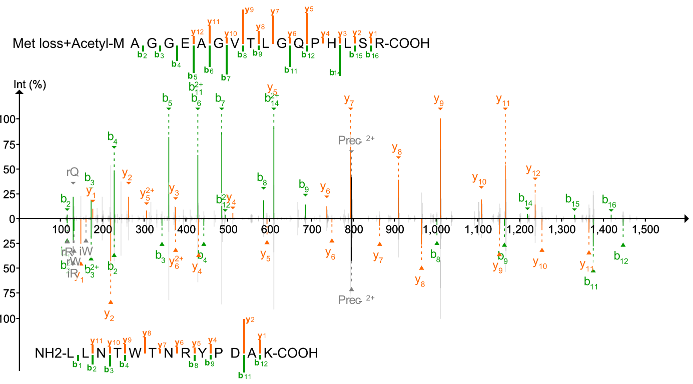


Scan 33273

Figure S7: **Peptide annotations for chimera spectra with localized modifications.** Mirror plots of two experimental spectra, annotated by the corresponding pairs of peptides discovered using CONGA at the 1% FDR level. Here we have localized the mass-modifications for the modified peptides. For scan 47880, we have identified the oxidation of the leading Methionine and for scan 33273, we have identified the loss of methionine and the acetylation of the subsequent n-terminal. The scan numbers relate to the representative HEK293 dataset.

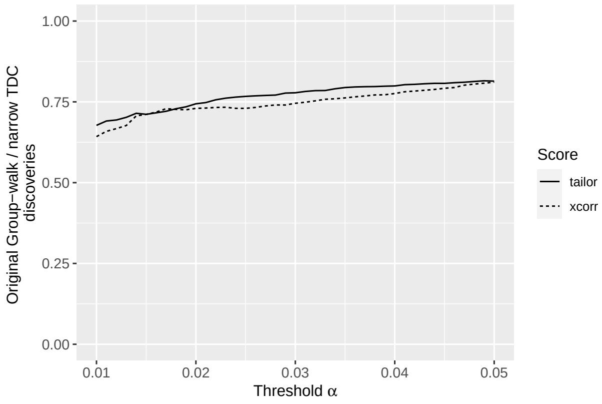


Figure S8: **The average ratio of Group-Walk to narrow-search TDC discoveries in real data.** The average ratio of the number of peptides discovered by Group-Walk to the corresponding number discovered by TDC. For TDC, the PSMs were generated by Tide using both XCorr and Tailor scores in narrow-search mode applied to one dataset (`PXD023571`) of our PRIDE-20 (Section 3; PRIDE-20 searches). The average was taken with respect to 20 randomly shuffled decoy databases. Notably, on average Group-walk loses over 25% of the peptides reported for this dataset by narrow-TDC at the 1% FDR level.

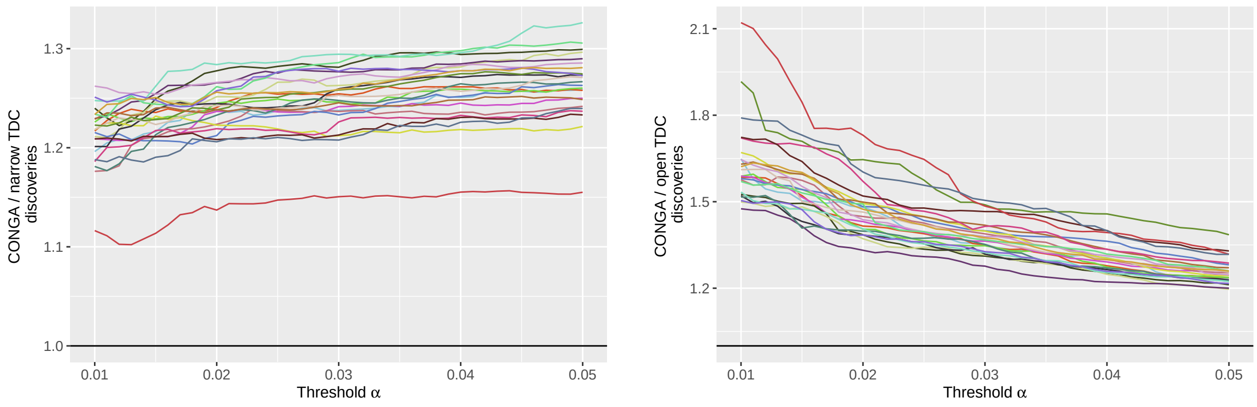


A B

Figure S9: **CONGA also improves on TDC when using localization-aware open search.** The figure plots the ratio between the number of peptides discovered by CONGA and the number of peptides discovered by TDC in the narrow-search (A), and in the open-search (B) for each of the 24 HEK293 spectrum files. Details regarding the search settings used are given in Section 3 (HEK293 searches), with the exception that we used the localized-aware scores in the open-search, and changed the lower precursor-window to the smallest allowable value of $-230$ Da. CONGA consistently outperforms TDC in the narrow and open-search, where the open-search uses the localization-aware scoring as discussed in [2].

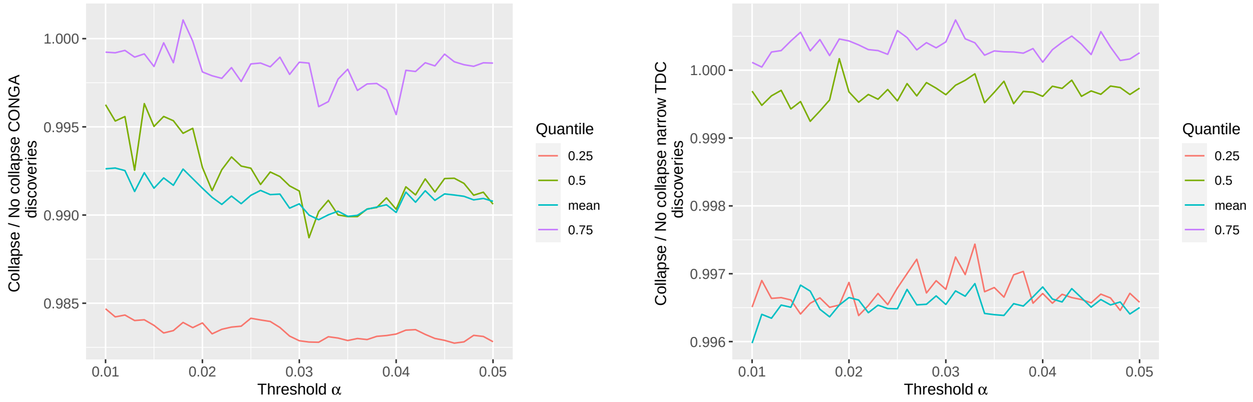


A B

Figure S10: **Comparing CONGA and narrow-TDC discoveries using peptide vs dynamic level analysis.** The figure plots the quartiles and mean of the average ratio between the number of peptides discovered by CONGA (A), and narrow-TDC (B) with and without collapsing to the representative sequence. The mean and quartiles are taken with respect to the 10 PRIDE-20 datasets that were deemed to contain variable modifications and the average of the number of representative-level to peptide-level discoveries is taken with respect to the 20 random decoys.
